# An efficient and accurate frailty model approach for genome-wide survival association analysis controlling for population structure and relatedness in large-scale biobanks

**DOI:** 10.1101/2020.10.31.358234

**Authors:** Rounak Dey, Wei Zhou, Tuomo Kiiskinen, Aki Havulinna, Amanda Elliott, Juha Karjalainen, Mitja Kurki, Ashley Qin, FinnGen, Seunggeun Lee, Aarno Palotie, Benjamin Neale, Mark Daly, Xihong Lin

## Abstract

With decades of electronic health records linked to genetic data, large biobanks provide unprecedented opportunities for systematically understanding the genetics of the natural history of complex diseases. Genome-wide survival association analysis can identify genetic variants associated with ages of onset, disease progression and lifespan. We developed an efficient and accurate frailty (random effects) model approach for genome-wide survival association analysis of censored time-to-event (TTE) phenotypes in large biobanks by accounting for both population structure and relatedness. Our method utilizes state-of-the-art optimization strategies to reduce the computational cost. The saddlepoint approximation is used to allow for analysis of heavily censored phenotypes (>90%) and low frequency variants (down to minor allele count 20). We demonstrated the performance of our method through extensive simulation studies and analysis of five TTE phenotypes, including lifespan, with heavy censoring rates (90.9% to 99.8%) on ~400,000 UK Biobank participants with white British ancestry and ~180,000 samples in FinnGen, respectively. We further performed genome-wide association analysis for 871 TTE phenotypes in UK Biobank and presented the genome-wide scale phenome-wide association (PheWAS) results with the PheWeb browser.

## Introduction

Survival models, especially the Cox proportional hazard model^1^, have been widely used to analyze time-to-event (TTE) outcomes, both in biomedical research^2–4^, and in genome-wide association studies (GWAS)^5–11^. It has been shown that the proportional hazard model can increase the power to detect genetic variants associated with the age-of-onset of TTE phenotypes in cohort studies compared to modelling the disease status using a logistic regression model^12–14^. With the availability of detailed time-stamped diagnosis data from Electronic Health Records (EHR), large biobanks, such as UK Biobank (UKBB)^15^ (> 400,000 individuals) and FinnGen (https://www.finngen.fi/en) (> 200,000 individuals), provide unprecedented opportunities to analyze TTE phenotypes to unravel the complex genetic architectures of disease onset, progression, and lifespan. Genome-wide scans of TTE phenotypes in large biobanks can potentially identify novel genetic variants associated with the onset of human diseases by leveraging both the disease status and the age-of-onset information.

In GWAS analysis, population structure and sample relatedness are often key factors that need to be controlled for. Biobank cohorts often have substantial population structure and relatedness. For example, in the UK Biobank, 91,392 out of 408,582 subjects with White British ancestry have at least one relative (up to 3^rd^ degree) in the data. Several linear and logistic mixed effects models have been developed to account for relatedness in GWASs for quantitative and binary phenotypes. To account for related subjects in the proportional hazard model, frailty models, which are mixed effects survival models, have been proposed^21,22^, where event times are assumed to be independent conditional on unobserved random effects called “frailties”. The frailties are modeled based on the dependence and clustering structure of the observations. Previous research has extensively studied shared frailty models with Gamma-distributed frailties^21,23–27^. However, the shared frailty model is limited in its scope to model more complicated dependency structures that arise in cohort-based association studies. To model complicated dependency structures, such as known familial structures and cryptic relatedness, the multivariate frailty model with Gaussian frailty was proposed^28,29^, and was later implemented in the R package COXME^30^, which, however, lacks scalability for GWASs. Recently the COXME method was further improved in COXMEG^31^, which utilizes several computational optimization strategies to make it applicable in genetic association studies, but COXMEG still cannot handle biobank-scale genome-wide datasets. Based on our performance benchmarking, even for 20,000 subjects, COXMEG requires 3,356 CPU-hours to perform a GWAS of 46 million variants, which means even with perfect parallelization on 30 CPUs, it would take over 4.6 days to complete the GWAS.

In large-scale GWASs, the score test is particularly useful among different asymptotic tests, because it requires fitting the model only once under the null hypothesis of no association^20^. Score tests have also been implemented in the COXMEG package^32^. However, score tests can lead to severe type I error inflation for phenotypes with heavy censoring, which is extremely common in biobank-based phenotypes. In the UK Biobank phenome that we built (see **ONLINE METHODS**), 871 TTE phenotypes have at least 500 events (cases), out of which 811 phenotypes have censoring rate more than 95%. The inaccuracies of the score test in unbalanced case-control phenotypes have been previously shown for logistic regression and logistic mixed effects models^19,33–35^, and a saddlepoint approximation^36^ (SPA)-based adjustment has been proposed and successfully implemented^19^ to accurately calibrate the p-values in such scenarios. Recently, the SPACox^11^ method also used SPA to calibrate p-values for time-to-event phenotypes in unrelated samples. However, the SPACox method does not account for sample-relatedness. Through simulations, we show similar inaccuracies are also present in score tests in frailty models for analyzing heavily censored phenotypes.

Here we propose a novel method for genome-wide survival analysis of TTE phenotypes, which accounts for both population structure and sample relatedness, controls type I error rates even for phenotypes with extremely heavy censoring, and is scalable for genome-wide scale PheWASs on biobank-scale data. Our method, Genetic Analysis of Time-to-Event phenotypes (GATE), transforms the likelihood of a multivariate Gaussian frailty model to a modified Poisson generalized linear mixed model (GLMM^20,37^) likelihood, employs several state-of-the-art optimization techniques to fit the modified GLMM under the null hypothesis, and then performs score tests calculated using the null model for each genetic variant. To obtain well-calibrated p-values for heavily censored phenotypes, GATE uses the SPA to estimate the null distribution of the score statistic instead of the traditionally used normal approximation. Moreover, our method saves the memory requirement substantially by storing the raw genotypes in binary format and calculating the elements of the GRM on the fly instead of storing or inverting a large dimensional GRM.

Through extensive simulations and analysis of TTE phenotypes from the UK Biobank data of 408,582 subjects with White British ancestry and the FinnGen study, we showed that GATE is scalable to biobank-scale GWASs of TTE phenotypes with type I error rates well controlled even for less frequent variants and heavily censored phenotypes. Benchmarking has shown that GATE can analyze 46 million variants in a GWAS with 408,582 subjects in ~ 14.5 hours using 30 CPUs with peak memory usage under 11 GB.

## Results

### Overview of Methods

GATE consists of two main steps: 1) Fitting the null frailty model to estimate the variance component and other model parameters, and 2) performing a score statistic-based test for association between each genetic variant and the phenotype. Step 1 involves iteratively fitting the null frailty model using similar optimization strategies as described in GMMAT^20^ and SAIGE^19^, such as using the computationally efficient average information restricted maximum likelihood (AI-REML^20,38^) algorithm for estimating the variance component, and using pre-conditioned gradient descent (PCG^39^) method to solve linear systems to avoid inverting the NxN genetic relatedness matrix (GRM). GATE computes the elements of the GRM on-the-fly when needed using binary vectors of raw genotypes, and thus it doesn’t require to supply, store, or invert a precomputed GRM, which can be extremely time and memory-consuming for large sample sizes (N). For example, in UK Biobank data with *M* = 93,511 markers and *N* = 408,582 subjects with White British ancestry, the memory requirement drops from 622 GB for storing a pre-computed GRM in floating point numbers, to only 8.9 GB for storing the raw genotypes in the binary format.

Step 2 involves scanning the entire genome and testing each variant for association using the score statistic. Since the overall cost of computing the variance of the score statistic for all variants is extremely high because it involves operations on the large-dimensional GRM, in step 2, GATE uses a variance ratio approximation commonly used in existing LMM and GLMM-based methods such as GRAMMAR-Gamma^17^, BOLT-LMM^16^, fastGWA^18^, and SAIGE^19^. The ratio of the variance of the score statistic with and without the random effects (and an attenuation factor due to estimating the baseline hazards) is computed using a subset of genetic markers. Previously, it was shown that this variance ratio remains approximately constant for variants with MAF ≥ 20 for LMM and GLMMs. Through analytical derivations and simulation examples, we show this observation to hold for frailty models as well (**Supplementary Note section 3 and Supplementary Figure 14**). Therefore, when performing the genome-wide scan, the variance of the score statistic is computed without using the GRM and then calibrated using the variance ratio.

Next, GATE uses the saddlepoint approximation^36^ (SPA) to approximate the null distribution of score statistics for association tests. SPA-based tests have been successfully used for logistic regression^34^ and logistic mixed models^19^ and provide more accurate p-values than traditional score tests under normal approximation for low-frequency variants when the case-control ratio is unbalanced. In GATE, we have implemented an efficient SPA-based test for frailty models that is similar to the fastSPA method in Dey et al.^34^. Through simulations and real data analysis, we show that SPA tests provide accurate and calibrated p-values, even for low-frequency variants when the censoring rate is high to 99%.

Both GATE and COXMEG^31^ conduct genetic association tests for TTE phenotypes using the frailty model. Besides the use of SPA-based tests, GATE uses the variance ratio approach to approximate the variances of the score statistics, while COXMEG calculates the variances using the GRM. Using simulation studies, we have shown that GATE provides consistent association p-values to COXMEG (*R*^2^ of −log10 P-values > 0.99) for common variants (MAF > 5%) when the censoring rate is 50% (**Supplementary Figure 1A**) and has well controlled type I error rates, even for less frequent variants and phenotypes with heavy censoring rates (**Supplementary Figure 1B**).

### Computation and Memory costs

To assess the computational performance of GATE and the score test implemented in the COXMEG package (COXMEG-Score), we randomly sampled subsets of different sample sizes from 408,582 UK Biobank subjects with White British ancestry. We then benchmarked association tests for overall lifespan (16,375 events, 389,721 censored) adjusting for the top four ancestry principal components, birth year and sex using GATE and COXMEG-Score on 200,000 variants randomly selected from 46 million genetic variants with imputation info ≥ 0.3 and MAC ≥ 20. In Step 1, 93,511 high-quality genotyped markers were used for the GRM. The projected overall computation time (**Figure 1 and Supplementary Table 1**) for GATE to analyze 46 million variants on *N* = 408,582 subjects was 318 CPU-hours, and the actual computation time on a machine with 30 cores was 14.5 hours. Step 2, which accounts for the majority of the computation time (95.4% for *N* = 408,582) requires substantially less memory (peak memory usage 0.85 GB) than Step 1 (peak memory usage 10.6 GB). However, even for 20,000 subjects, the projected computation time and memory usage for COXMEG-Score were 3,356 CPU-hours and 32.75 GB, compared to only 34 CPU-hours and 0.74 GB required by GATE, achieving 99% and 97.7% reductions in computation time and memory, respectively. This means even with perfect parallelization on 30 CPUs, COXMEG-Score would require 4.6 days to complete the GWAS with only 20,000 subjects. The observations also suggest that the computation time and memory requirements increase nearly linearly with the sample size for GATE, whereas they increase quadratically for COXMEG-Score.

**Figure 1:**
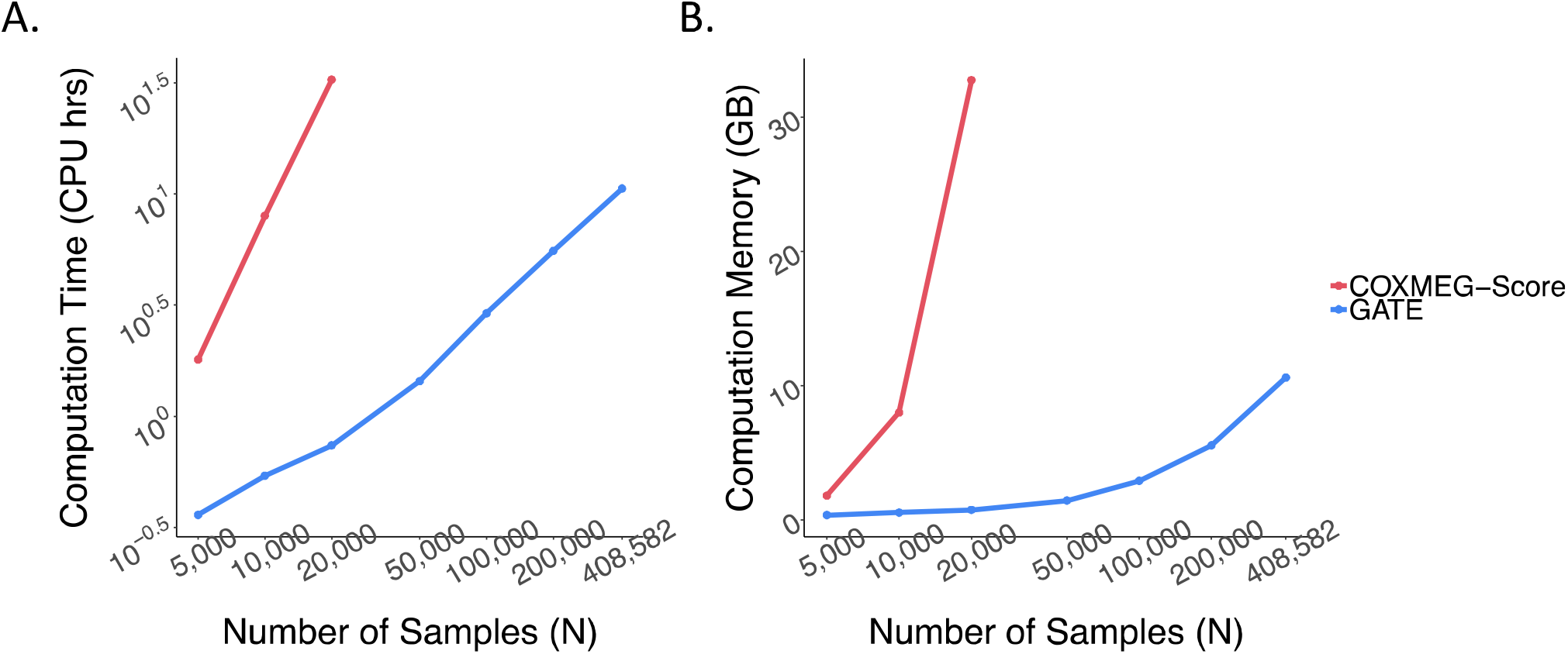
Projected computation time (A) and memory usage (B) for GATE and COXMEG-Score as a function of sample size (N). The numerical data are provided in Supplementary Table 1. Benchmarking was performed for the GWAS of lifespan based on randomly subsampled data from UK Biobank White British ancestry subjects. Association tests were performed on 200,000 randomly selected markers with imputation INFO ≥ 0.3, with the filtering criteria of MAC ≥ 20. The computation times were projected for testing 46 million variants with INFO ≥ 0.3 and MAC ≥ 20. The reported run times are medians of five runs, each with randomly sampled subjects with different randomization seeds. The x and y axes are plotted in log10 scale.

### Phenome-wide GWAS of time-to-event phenotypes in the UK Biobank data

We have applied GATE to perform phenome-wide GWAS for 871 UKBB TTE phenotypes with at least 500 events, adjusting for top four PCs, birth year, and sex (except for 93 sex-specific phenotypes). The TTE phenotypes were created based on the International Classification of Disease (ICD) codes version 9 and 10 mapped to the PheWAS code (PheCode^40^) definitions (See **ONLINE METHODS**) as well as their associated diagnosis dates in the UK Biobank electronic medical records. For each phenotype, we analyzed approximately 46 million genetic markers imputed from the Haplotype Reference Consortium^41^ panel and UK10K^42^ with imputation INFO score ≥ 0.3 and MAC ≥ 20. Among the 408,582 UK Biobank subjects with White British ancestry, 91,392 had at least one relative up to third degree^15^. To account for the relatedness among the subjects, we used 93,511 high-quality genotyped markers with MAF ≥ 0.01 to construct the GRM in Step 1. The same set of markers were used by the UK Biobank research group^15^ for estimating kinship among the samples because they are only weakly informative of the ancestry and therefore provide more accurate kinship estimates. We also performed a sensitivity analysis using a larger set of markers (245,745) for the four exemplary phenotypes discussed before (See **Supplementary Note Section 7**). We further applied SPA-based adjustment of the score test because to the censoring rates (**Supplementary Figure 2**) were extremely high for most of the TTE phenotypes in the UKBB (for example, 811 out of 871 have censoring rate more than 95%). The summary statistics for all 871 PheCodes analyzed using GATE are available to download from a public repository (see URL) and browsed in the PheWeb^43^ (see URL).

Here we discuss the association results using four phenotypes with different censoring rates as exemplars: ischemic heart disease (IHD: PheCode 411, N events=36,962, N censored=370,814, censoring rate=90.9%), female breast cancer (FBC, PheCode 174.1, N events=15,396, N censored=192,764, censoring rate=92.6%), glaucoma (PheCode 365, N events=6,046, N censored=392,925, censoring rate=98.5%), and Alzheimer’s Disease (AD: PheCode 290.11, N events=822, N censored=342,059, censoring rate=99.8%). The Manhattan and QQ plots for the GWAS of these phenotypes using GATE with and without SPA are presented in **Figure 2** and **Figure 3**, respectively. The results demonstrate that not adjusting for SPA greatly inflates the type I errors, especially for the low frequency variants, whereas the SPA-adjusted method shows well controlled type I error rates. In total, 114 loci have been identified for the four TTE phenotypes: 55 for IHD, 37 for FBC, 19 for glaucoma, and 3 for AD. We also applied GATE to these four phenotypes in the FinnGen study (see **ONLINE METHODS**) and 81 out of the 114 loci were also tested in the FinnGen study, of which 78 had the same effect direction in both UKBB and FinnGen. 69 out of the 81 loci were successfully replicated in FinnGen with p-value < 0.05. The complete list of all significant loci and the association results in the UKBB, FinnGen as well as the meta-analysis of the two data sets are reported in **Supplementary Table 2**. Overall, 99 out of the 114 significant loci have been previously reported to be associated with disease risk in case-control studies to the best of our knowledge. Several loci that are previously well known as associated with the risk of the diseases have been identified in our study. For example, the loci *LPA* and *CELSR2* for IHD^44^,^45^, *FGFR2^4^* and *CASC16^47^* for breast cancer, *MYOC*^48^ and *TMCO1^49^* for glaucoma, and *APOE* e4 variant for AD^50^. The age-varying predicted risk of disease onset based on the GATE method, and the age-varying disease-free probability by genotypes based on the Kaplan-Meier curve^51^ for the exemplary top hits were plotted in **Figure 4** and **Supplementary Figure 3**, respectively.

**Figure 2:**
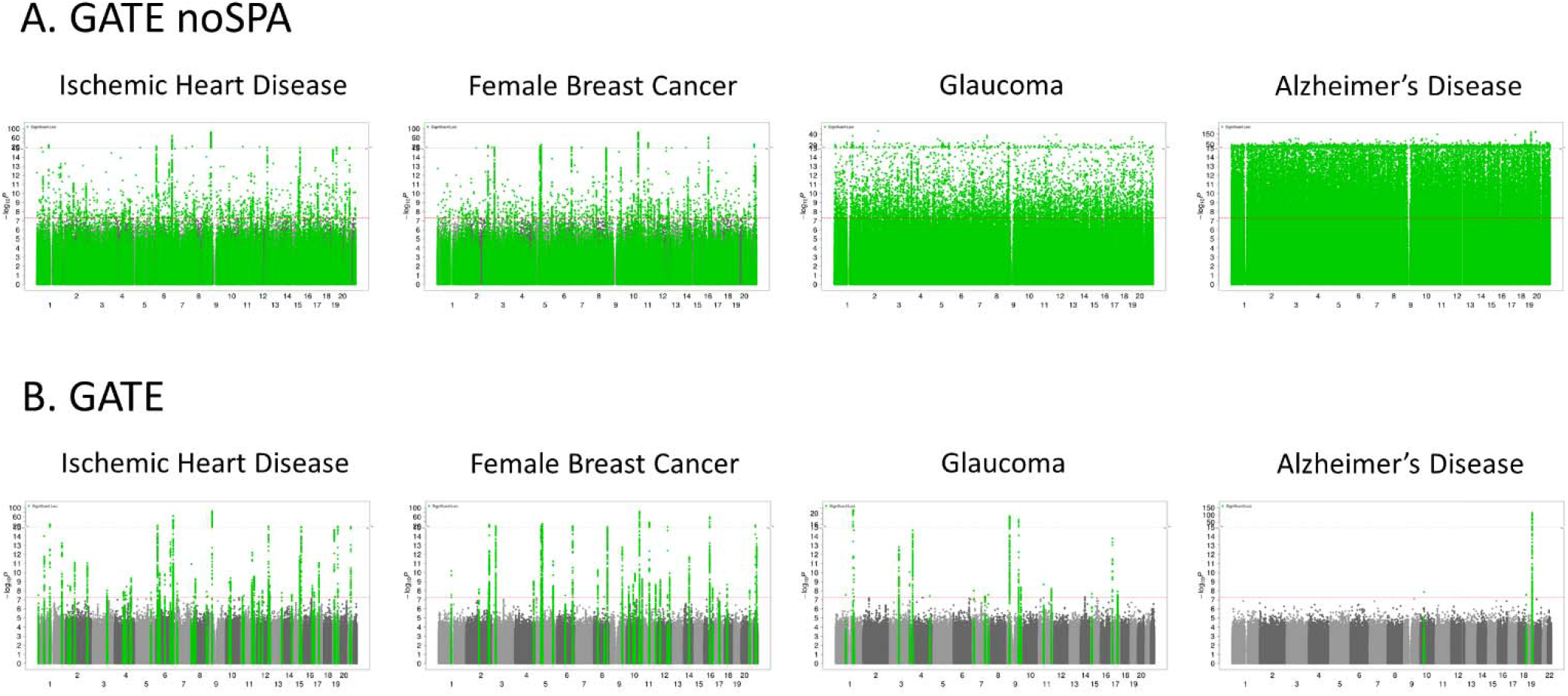
Manhattan plots for GWAS of four time-to-event phenotypes with different censoring rates in the UK Biobank data with White British ancestry: GWAS results using GATE-noSPA (**A)** and GATE (B) are shown for ischemic heart disease (PheCode 411, N=407776, censoring rate=90.9%), female breast Cancer (PheCode 174.1, N=208160, censoring rate=92.6%), glaucoma (PheCode 365, N=398971, censoring rate=98.5%), and Alzheimer’s Disease (PheCode 290.11, N=342881, censoring rate=99.8%).

**Figure 3:**
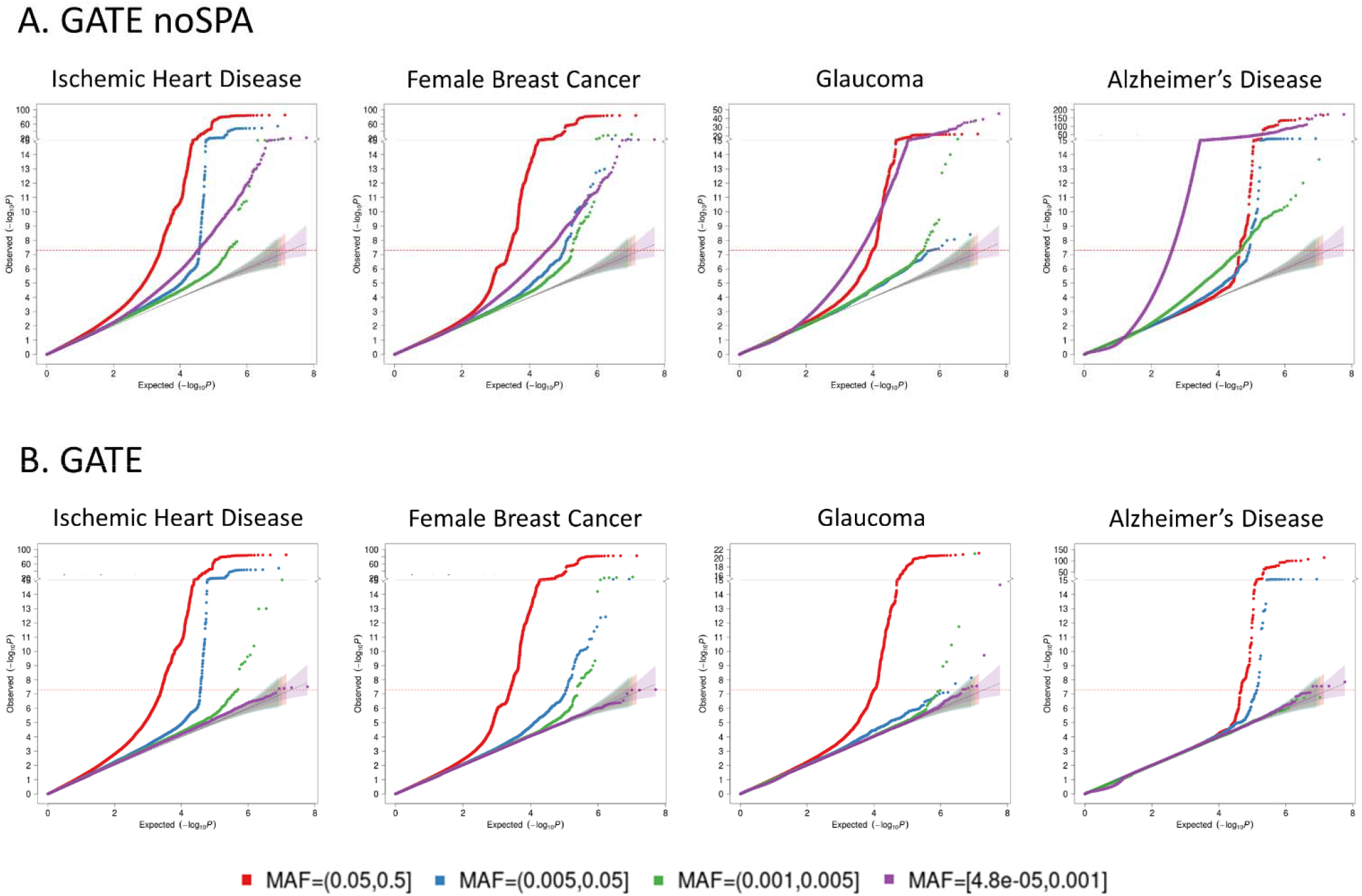
Quantile-quantile (QQ) plots for GWAS of four time-to-event phenotypes with different censoring rates in the UK Biobank data with White British ancestry: GWAS results using GATE-noSPA (A) and GATE (B) are shown for ischemic heart disease (PheCode 411, N=407776, censoring rate=90.9%), female breast Cancer (PheCode 174.1, N=208160, censoring rate=92.6%), glaucoma (PheCode 365, N=398971, censoring rate=98.5%), and Alzheimer’s Disease (PheCode 290.11, N=342881, censoring rate=99.8%). QQ plots are color-coded based on different minor allele frequency categories.

**Figure 4:**
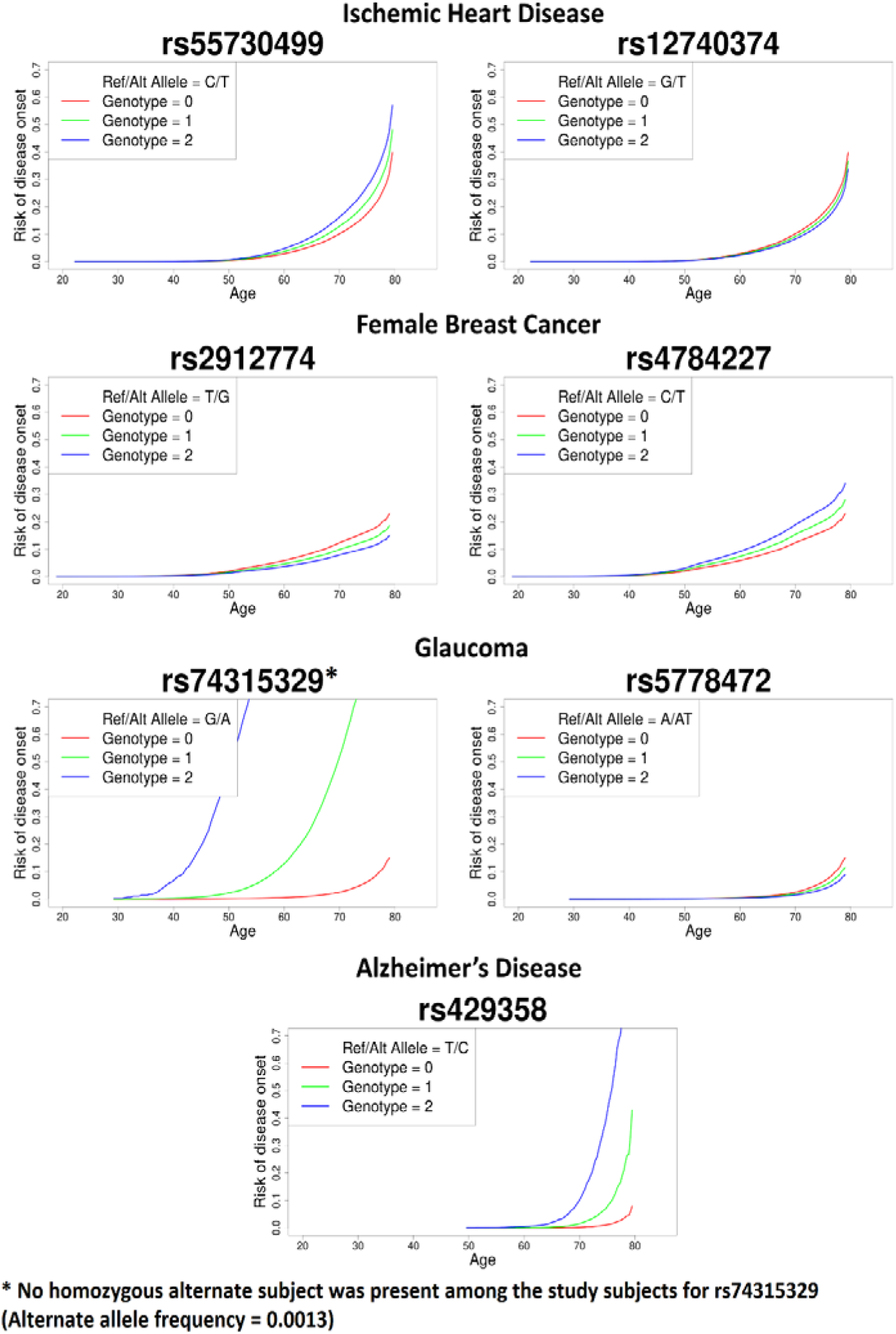
Predicted risk of disease onset over-time by genotypes for loci LPA and CELSR2 for ischemic heart disease, FGFR2 and CASC16 for female breast cancer, MYOC and TMCO1 for glaucoma, and APOE e4 variant for AD. The red, green and blue lines represent the risk of disease onset for alternate allele counts zero, one and two, respectively for a female subject born in 1950 (median birth year in the UKBB data) with the top four PC coordinates each set at the mean level across the UK Biobank subjects with white British ancestry.

### GWAS of lifespan in the FinnGen Study and the UK Biobank

We have also applied GATE to the overall lifespan in the FinnGen study (N events = 15,152, N censored = 203,244), in which the age of death ranges from 7 years old to 106 years old as shown in **Supplementary Figure 4**. We identified the previously reported *APOE* locus for lifespan^52^ in FinnGen, in which the most significant variant is the APOE-e4 missense variant rs429358 (MAF = 18.3%, p-value = 1.01 x 10^-14^) and it is well-known to be associated with lifespan, cardiovascular diseases, stroke, and Alzheimer’s disease^53–55^. This locus has been replicated in UKBB (N events = 16,375 and N censored = 389,721, see **Supplementary Figure 5**) with p-value 1.92 x 10^-5^ and meta-analysis p-value 4.04 x 10^-17^ (**Supplementary Table 3** and **Supplementary Figure 6**). The top hit in UKBB (rs157592, MAF = 18.7%, p-value = 1.87 x 10^-8^) had LD r^2^ = 0.7 with rs429358 as presented in the **Supplementary Table 3.** This variant is in the intergenic region and have no in-silico functions according to the FAVOR functional annotation online portal^56^ (See URL).

### Simulation Studies

We investigated the type I error rates and power of GATE in presence of sample relatedness using 10,000 simulated samples. Due to computation burden, we used GATE-noSPA instead of COXMEG-Score for type I error evaluation as **Supplementary Figure 1C** shows the two approaches provide consistent association p-values (*R*^2^ of −log10 p-values > 0.99).

The type I error rates of GATE were evaluated based on association tests of 9.4×10^8^ simulated genetic markers on 10,000 samples, which contain 500 families and 5,000 independent samples. Each family has 10 members, simulated based on the pedigree shown in **Supplementary Figure 7**. The variance component parameter τ is set to be 0.1 and 0.25 (see **ONLINE METHODS**). The empirical type I error rates at the significance level α = 1×10^−6^ and 5×10^−8^ are shown in the **Supplementary Table 4 and Supplementary Figure 8A.** Our simulation results suggest that GATE has well controlled type I error rates even for low frequency variants (down to MAC = 20) when the phenotype is heavily censored (90%). However, without SPA, the score tests in GATE suffer from inflated type I error rates as the case-control ratios become more unbalanced and the frequency of variants decreases. We also evaluated type I error rates of GATE in a setting with cryptic sample relatedness by randomly selecting 10,000 UKBB participants with white British ancestry. Phenotypes were simulated using the real genotypes to mimic the sample relatedness of a real-world dataset, and association tests were conducted on the imputed genetic markers in the UKBB (see **ONLINE METHODS**). Similarly, we observed that the type I error rates were well controlled in GATE in presence of cryptic sample relatedness with different censoring rates (**Supplementary Table 5, Supplementary Figure 8B and 9**).

Next, we evaluated empirical power of GATE at α = 5×10^−8^ and compared to the power of COXMEG-Score. **Supplementary Figure 10** shows the power curve by hazard ratios for variants with MAF 0.05 and 0.2 when τ =0.25 and the censoring rate = 50%. Both methods have nearly identical power in all simulation settings. We do not compare their powers in the presence of heavy censoring, in view of the inflated type I error rate of COXMEG-Score.

Overall simulation studies show that GATE can control type I error rates even when censoring rate is high and has similar power for common variants as COXMEG-Score. In contrast, same as GATE-noSPA, COXMEG suffers type I error inflation and the inflation is especially severe with low MAF and heavy censoring (**Supplementary Figure 1B, 1C, 8** and **9**).

## Discussion

In this paper, we have proposed a novel method to perform scalable genome-wide survival association analysis of censored TTE phenotypes in large biobanks using an efficient implementation of the frailty model. Our method can adjust for population structure and sample relatedness and provide accurate p-values even in extreme cases of very low frequency variants and heavily censored phenotypes (incidence rate < 0.1%). Applying this approach to the UK Biobank and the FinnGen study, we demonstrated that our method is scalable to the analysis of large biobank-scale datasets with > 400,000 subjects.

Biobanks with genetic data linked to EHR records/survey questionnaires provide unprecedented opportunities for genetic association studies on TTE phenotypes to identify genetic risk factors that affect the onset and progression of diseases. However, biobanks pose challenges to such analysis because of the high computational and memory cost required to handle large data sets with extensive population structure and relatedness. Moreover, current methods artificially inflate associations when heavily censored phenotypes (e.g., censoring rate > 75%) and low frequency variants (MAF < 1%) are involved. The proposed method, GATE performs a frailty model-based association analysis to account for both population structure and relatedness using score tests with SPA adjustment, which provides accurate p-values under heavy censoring. In addition, it implements several optimization techniques that were previously used in the context of linear and logistic mixed models in BOLT-LMM and SAIGE to make it computationally feasible to analyze large biobank cohorts. We have applied GATE to 871 TTE phenotypes in the UK Biobank data with White British ancestry, which were constructed based on PheCodes mapped to ICD codes and have at least 500 events. The genome-side summary statistics are available for public to download. We have also created a PheWeb^43^ for users to explore and visualize the PheWAS results.

TTE phenotypes are particularly suited not only for studying disease onsets, but also for exploring other progression phenotypes such as times of surgery, recurrence, times of onset of secondary phenotypes after an initial diagnosis etc. Previously, the lack of scalable GWAS methods for TTE outcomes has hindered such investigations in massive scales. By facilitating large-scale GWAS of TTE phenotypes, GATE opens the door to such deeper investigations.

One consideration while analyzing TTE phenotypes is the appropriate choice of the unit of time. To assess the impact of time-units on the GWAS results, we performed sensitivity analysis using the event and censoring times rounded to the nearest 1 month, 3 months, 6 months and 12 month time-units for the four exemplary UK Biobank phenotypes presented in this paper, and compared the p-values across different time-units (**Supplementary Figure 11**). The p-values were very similar across the four time-units for all phenotypes, with more detailed time-units resulting in slightly more significant p-values.

For the selection of number of markers to construct the GRM, there is a trade-off between computation cost and the accuracy of adjusting the sample relatedness. Increasing the number of markers (*M)* included in the GRM linearly increases the computation time and memory requirement of step 1, whereas using too few markers may not be sufficient to capture the detailed familial and cryptic relatedness among the samples properly^57^. For the UK Biobank data analysis, we used *M* = 93,511 LD pruned high-quality genotyped markers which were used by the UK Biobank research group for estimating kinship among the samples^15^. We performed a sensitivity analysis (see **Supplementary Note Section 7**) by increasing the number of markers to *M* = 245,975 pruned markers with MAF ≥ 0.01. The results (**Supplementary Figure 12 and 13**) showed that the p-values were generally concordant, and the p-values using *M* = 245,975 markers were slightly larger than the p-values using *M* = 93,511 markers.

There are several limitations to GATE. First, similar to other mixed model methods for genetic association tests, the computation time required for the algorithms to converge in step 1 can vary among different phenotypes and study samples because of the difference in heritability and the extent of sample relatedness. Second, GATE uses a score statistic-based test without fitting the model under the alternate hypothesis, which can be computationally inefficient. Therefore, it does not provide accurate estimates of hazard ratios for the genetic variants. Following a similar approach as in several other mixed model-based methods^16,17,19,58^, GATE provides a hazard ratio estimate using the null model parameter estimates (see **Supplementary Note Section 5**). Third, the current implementation of GATE is targeted to perform single-variant association analysis, which can suffer from low power to detect associations in extremely rare variants. With whole genome and whole exome sequencing data available, a possible future extension of this method can allow for mask-based or region-based association tests to improve power for the rare variants^56,59^. Finally, the current version of GATE does not incorporate left-truncated data, which may not be valid for early-onset phenotypes in biobanks with relatively older participants. For example, the median age of UK Biobank’s participants is 59 years old and the earliest dates of health data available are around late 1990s, and assuming no left-censoring can reduce association power for early-onset diseases. The next work will extend GATE to allows for left-truncated phenotypes. In summary, we have proposed a scalable and accurate method, GATE, to perform genome-wide PheWAS of TTE phenotypes on large biobank cohorts accounting for population structure, sample relatedness and heavy censoring. We demonstrated that it is possible to efficiently analyze the current largest biobank (UK Biobank) of > 400,000 subjects using GATE. Our method facilitates biobank-based PheWAS of TTE phenotypes which ultimately contributes towards identifying genetic components that affect the onset and progression of complex diseases.

## URLs

GATE is implemented as an open-source R package available at https://github.com/weizhou0/GATE. The GWAS results for 871 time-to-event phenotypes in UK Biobank using GATE are currently available for public download at http://gate.genohub.org/. Manhattan plots, Q-Q plots, and regional association plots for each TTE phenotype as well as the PheWAS plots can be browsed at http://phewas.genohub.org/. The FAVOR^56^ portal is accessed through favor.genohub.org.

### Acknowledgments

The FinnGen project is funded by two grants from Business Finland (HUS 4685/31/2016 and UH 4386/31/2016) and eleven industry partners (AbbVie Inc, AstraZeneca UK Ltd, Biogen MA Inc, Celgene Corporation, Celgene International II Sàrl, Genentech Inc, Merck Sharp & Dohme Corp, Pfizer Inc., GlaxoSmithKline, Sanofi, Maze Therapeutics Inc., Janssen Biotech Inc). Following biobanks are acknowledged for collecting the FinnGen project samples: Auria Biobank (www.auria.fi/biopankki), THL Biobank (www.thl.fi/biobank), Helsinki Biobank (www.helsinginbiopankki.fi), Biobank Borealis of Northern Finland (https://www.ppshp.fi/Tutkimus-ja-opetus/Biopankki/Pages/Biobank-Borealis-briefly-in-English.aspx), Finnish Clinical Biobank Tampere (www.tays.fi/en-US/Research_and_development/Finnish_Clinical_Biobank_Tampere), Biobank of Eastern Finland (www.ita-suomenbiopankki.fi/en), Central Finland Biobank (www.ksshp.fi/fi-FI/Potilaalle/Biopankki), Finnish Red Cross Blood Service Biobank (www.veripalvelu.fi/verenluovutus/biopankkitoiminta) and Terveystalo Biobank (www.terveystalo.com/fi/Yritystietoa/Terveystalo-Biopankki/Biopankki/). All Finnish Biobanks are members of BBMRI.fi infrastructure (www.bbmri.fi). This research has been conducted using the UK Biobank Resource under application number 52008. X.L. was supported by NCI R35-CA197449, P01-CA134294, U19-CA203654 and NHLBI R01-HL113338. B.M.N. was supported by NHGRI U01-HG009088-04S3 and NIMH R37-MH107649-06. R.D. was supported by NCI R35-CA197449. W.Z._was supported by an NIH T32 fellowship (Grant number: 1T32HG010464-01). A.P. was supported by the Academy of Finland Centre of Excellence in Complex Disease Genetics (Grant No. 312074)

### Author Contributions

R.D., W.Z. X.L., B.M.N., and M.J.D. designed experiments. R.D. and W.Z. performed experiments. R.D. and W.Z. implemented the software with input from X.L., B.M.N., and M.J.D. R.D. constructed phenotypes for UK Biobank data. R.D. and X.L. analyzed UK Biobank data. A.Q., R.D., and W.Z. created the PheWeb browser for UK Biobank results. W.Z., T.K., A.H., A.E., J.K., M.K., and A.P. analyzed data for the FinnGen study. Helpful advice was provided by S.L. R.D. and W.Z. wrote the manuscript with input from all co-authors.

### Online Methods

#### Frailty model for Time-to-event phenotypes

Consider a study of *N* subjects, where for the *i*-th subject, we observe the data pair (*δ_i_,t_i_*), where *δ_i_* is a censoring indicator, with *δ_i_* = 1 if the *i*-th subject experiences an event during the study period, and *δ_i_* = 0 otherwise, i.e., censored. Let *t_i_* denote the observed event or censoring time. For the *i*-th subject, let the *p* x 1 vector *X_i_* denote the covariates, and *G_i_* = 0,1,2 denote the minor allele counts for the genetic variant of interest. Then, in a frailty model^25,28,60^, the conditional hazard function of subject *i* at time t given the covariates, genotype and random effect/frailty *b_i_* is modeled as

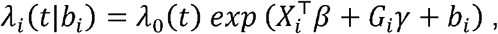

where *β* and *γ* are the regression coefficients of the covariates *X_i_* and the genotype *G_i_* respectively, and *Λ*_0_(*t*) is the baseline hazard function at time *t*, the frailty *b* ; (*b*_1_,…,*b_N_*) follows a multivariate normal distribution *N*(0,*τV*), with *V* being the Genetic Related Matrix (GRM). Unlike standard generalized linear mixed models, the covariate vector *X_i_* in a frailty model does not include the intercept term, instead the baseline hazard *Λ*_0_(*t*) works as the intercept in a frailty model. We test the null hypothesis of no genetic association *H*_0_:*γ* = 0 vs *H*_1_: *γ* ≠ 0.

#### Estimating the variance component and other null model parameters (step 1)

First, the likelihood for the observed event status-time pairs (*δ_i_*, *t_i_*) under the frailty model is derived and expressed as a modified Poisson mixed effects model likelihood, with the mean function weighted by the cumulative baseline hazard (CBH) function 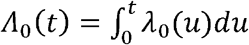. The CBH function is estimated by the Breslow’s estimator 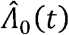 as a step function. Breslow showed that the maximum likelihood approach for the proportional hazard model (for unrelated subjects) that leads to the estimator 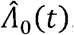, is equivalent to maximizing the partial likelihood proposed by Cox^1^. In the **Supplementary Note Section 6**, we have shown that the same maximum likelihood approach holds for frailty models (related subjects) as well given the random effects. Then, using the penalized quasi-likelihood (PQL^37^) method and the AI-REML^38^ algorithm, the model parameters under *H_o_* are estimated iteratively. To avoid storing large *N* × *N* GRMs, GATE only calculates the elements of the GRM when they are needed using raw binary format genotypes. For scalable computation of quantities of the form *A*^−1^ *x* that arises in the model fitting steps, where *A* is a large matrix and *x* is a vector, GATE uses the PCG algorithm^39^, which has been previously used in BOLT-LMM^16^ and SAIGE^19^ to accurately compute quantities like *y* = *A*^−^ *x* by solving the linear system of equations *Ay* = *x*, instead of explicitly inverting the large matrix *A*.

Once the null model parameters, random effects and cumulative baseline hazard functions 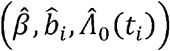 have been estimated, GATE estimates the variance ratio from a small number of markers. Denote the fitted means by 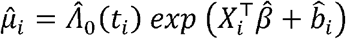, and the weight matrix 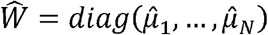. Then the score statistic, under *H*_0_:*γ* = 0 is 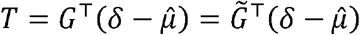, where 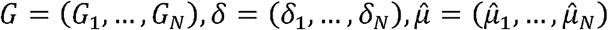. The covariate-and-intercept-adjusted genotypes are denoted by 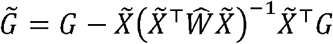, where 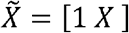 is the augmented covariate matrix. Then, the variance of the score statistic under *H*_0_ is given by 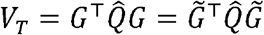, where 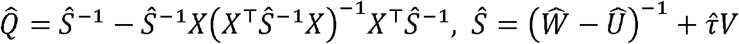. The expression of 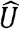 is described in detail in the **Supplementary Note Section 1.3**. Unlike in the GLMMs, the term 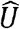 appears in the variance of the score statistic due to the attenuation of information (additional variability) for estimating *Λ*_0_(*t_i_*)s. The variance ratio is then calculated as 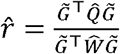 calculates the variance ratio based on 30 randomly selected genotyped markers with MAC ≥ 20 and computes the coefficient of variation (CV). If the CV of the variance ratios is smaller than 0.001, then the mean of the variance ratios is selected as 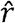, otherwise more markers are selected at an increment of 10 markers, and the CV is recalculated until the CV becomes smaller than 0.001.

#### Score test using SPA

Using the estimated variance ratio 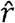, the variance-adjusted test statistic can be calculated as 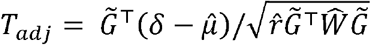, which under the null hypothesis has mean zero and variance unity. The traditional score test then assumes asymptotic normality of the score statistic *T* (and thus *T_adj_* as well) under *H*_0_, to calculate the p-value. However, observations have been made before in the context of logistic mixed models that the asymptotic normality assumption of the score test statistic leads to severe Type I error inflation for low-frequency and rare variants when the case-control ratio is unbalanced^19^. We make the same observations in frailty models as well when the censoring rate is high. In order to provide well calibrated p-values in such situations, we used saddle point approximation (SPA) to approximate the null distribution of the score statistic, which has been shown to have better approximation error bounds compared to the normal approximation^34,36,62,63^, especially at the extremely small tail probability region of *α* = 5 x 10^−8^. Contrary to the normal approximation which only utilizes the first two moments only to approximate, SPA utilizes the entire moment generating function (MGF). In fact, it uses the cumulant generating function (CGF), i.e., is the logarithm of the MGF, which for the frailty model, based on the modified Poisson mixed model likelihood, can be derived as 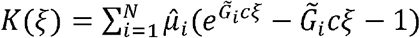, where 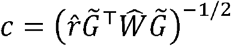. Then, the distribution of *T_adj_* can be calculated based on the SPA by 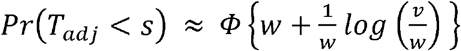, and the p-value is given by *p* = *Pr*(*T_adj_* < −|*s*|) + *Pr*(*T_adj_* > |*s*|), where *T_adj_* = *s* is the observed adjusted score statistic, 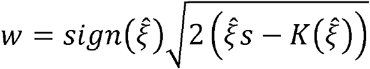, 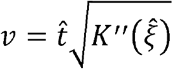, 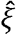 is the solution to the equation 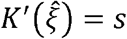, and 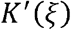 and 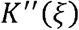 are the first and second derivatives of the CGF *K*(*ξ*), respectively.

Since the normal approximation works well around the mean, we use the normal approximation when *T_adj_* is less than two standard deviations away from the mean for faster computation. In addition, a faster version of the SPA similar to Dey et al.^34^ is also implemented which reduces the computation time even further, from *O*(*N*) to *O*(*N_c_*), where *N_c_* is the number of minor allele carriers.

#### Data Simulation

We carried out a series of simulations to evaluate the performance of GATE, including the type I error rates and power. To evaluate whether GATE can control type I error rates in presence of sample relatedness, we randomly simulated a set of 1,000,000 base-pair “pseudo” sequences, in which variants are independent to each other. Alleles for each variant were randomly drawn from Binomial(n = 2, p = MAF). Then we performed the gene-dropping^64^ simulation using these sequences as founder haplotypes that were propagated through the pedigree of 10 family members shown in **Supplementary Figure 7**. We simulated genotypes of 150,000 genetic variants with MAF ≥ 1% for 5,000 independent samples and 500 families based on the pedigree to estimate the GRM on-the-fly in Step 1 of GATE and genotypes of 1.9 million genetic variants with MAC ≥ 20 for association tests in Step 2. MAFs were randomly sampled from the MAF spectrum in UK Biobank imputation data as shown in **Supplementary Figure 9**. For each subject *i,* the censoring time *T_ci_* was randomly selected from exponential distribution with mean 1/*λ_c_* and the underlying failure time *T_fi_* was generated from a frailty model with the underlying exponential hazard function 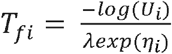, where *U_i_* ~ uniform (0,1) and *η_i_* is the linear predictor. Under the null hypothesis of no genetic effects, 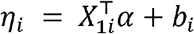, where *X*_1_ is a covariate that was randomly drawn from *N*(0,1), *α* is the coefficient and is 0.5 and *b_i_* is the random effect simulated from *N*(0,*τψ*) with *τ* = 0.1 and 0.25, respectively, which is the variance component parameter. The time for subject *i* is *t_i_* = *min*(*T_ci_, T_fi_*) and *δ_i_* ; *I*(*T_fi_* ≤ *T_ci_*). We selected *λ*, the mean of the exponential hazard function, corresponding to different censoring rates 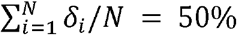, 75% and 90%. We repeated the simulation for 500 times. For each phenotype set, a null frailty model was fitted in Step 1 with the covariate *X*_1_. In Step 2, we conducted single variant association tests on 1.9 million simulated genetic markers. In totally, about 9.4×10^8^ association tests were conducted. We evaluated the empirical type I error rates at the type I error rate α = 1×10^−6^ and 5×10^−8^ as shown in **Supplementary Table 4** and **Supplementary Figure 8A**. These results have indicated that GATE can produce well calibrated type I error rates in the presence of sample relatedness at the significance level, while GATE-no SPA (similar to COXMEG) has inflated type I error rates and inflation gets larger than censoring rates is higher (**Supplementary Table 4**). For example, GATE-no SPA has type I error rate 8.9×10^−6^ at α = 5×10^−8^ when censoring rate is 75% and 2.8×10^−5^ when censoring rate is 90% with τ = 0.1.

To evaluate whether GATE can control type I error rates in presence of cryptic sample relatedness, we have randomly selected *N* = 10,000 samples with white British ancestry from UK Biobank and simulated TTE phenotypes based on the observed genotyped of these subjects in the approach described above for pedigree-based data sets, except that under the null hypothesis of no genetic effects, 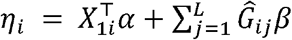 and was simulated based on real genotypes of randomly selected L = 30,000 LD-pruned (r2 < 0.2) markers from the odd chromosomes with MAF ≥ 1%. The real genotypes were used for simulating real sample relatedness in the null model. In particular, *X*_1_ is a covariate that was randomly drawn from *N*(0,1), *α* is the coefficient and is 1, 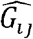 is the standardized genotype value for the jth marker of ith subject and *β* is the genetic effect size following *N*(0,*τ/L*). where *τ* = 0.25, which is the variance component parameter. The time for subject *i* is *t_i_* = *min*(*T_fi_*, ≥ *T_ci_*) and *δ_i_* = *T_ci_*). We selected *λ*, the mean of the exponential hazard function, corresponding to different censoring rates 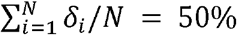, 75% and 90%. We repeated the simulation for 100 times. For each phenotype set, a null frailty model was fitted in Step 1 with covariates including the first 4 genetic principal components, which were estimated for all White-British participants in the UK Biobank, and *X*_1_. In Step 2, we conducted single variant association tests on genetic markers on the even chromosome. In totally, 8.3×10^8^ were conducted. We evaluated the empirical type I error rates at the type I error rate α = 1×10 and 5×10 as shown in **Supplementary Table 5** and **Supplementary Figure 8B**, which suggests that GATE produces well calibrated type I error rates in the presence of cryptic relatedness at the corresponding significance levels.

To evaluate the empirical power of GATE and compare the power to COXMEG, phenotypes were generated under the alternative hypothesis for 10,000 samples, which contain 500 families and 5,000 independent samples. The family pedigree is shown in the **Supplementary Figure 7**. We simulated 100 datasets with 10 genetic markers with different hazard ratios. Power was evaluated at α=5×10^−8^ with the censoring rate 50% for MAF 0.05 and 0.2 as presented in the **Supplementary Figure 10.**

#### Building the UK Biobank TTE Phenome

The time-to-event phenotypes for the UK Biobank were constructed as the disease phenotypes defined based on the hierarchical PheCodes^40^ that represent different disease groups. The ICD9 and ICD10 codes were mapped to PheCodes using a combination of available maps through the Unified Medical Language System (see URLs) and other sources, string matching, and manual review^19,40^. For each PheCode, the subjects who had the PheCode were regarded as having events, and the subjects who did not have the PheCode were regarded as censored. For each failed subject, the TTE (failure time) was calculated by subtracting the birth year from the earliest time of diagnosis of any of the PheCode-specific ICD codes, rounded to the nearest full month. To obtain the TTE (censoring time) for each censored subject, the birth year was subtracted from the time of the last non-imaging visit to any of the UK Biobank ascertainment centers, or the last time any ICD code was recorded for that subject, or the time of death if death was recorded during the course of the study, whichever is latest, rounded to the nearest full month. For lifespan, the subjects who had their death recorded, were assigned the failed status with the ages at death as the corresponding TTE, and the subjects who did not have their death recorded were assigned the censored status with the TTE defined as before.

#### FinnGen

FinnGen is a public-private partnership project combining genotype data from Finnish biobanks and digital health record data from Finnish health registries (https://www.finngen.fi/en). Release 5 analysis contains 218,792 samples after quality control with population outliers excluded via principal component analysis based on genetic data. TTE phenotypes were constructed from population registries and ICD10 codes, and harmonizing definitions over ICD8 and ICD9, including ischemic heart disease (N events=30,952, N censored=187838, censoring rate=85.8%), female breast cancer (N events=8,401, N censored=114,878, censoring rate=93.2%), glaucoma (N events=8,591, N censored=210199, censoring rate=96.1%) and Alzheimer’s disease (N events=3,899, N censored = 207,324, censoring rate=98.2%). We conducted genome-wide survival analysis using GATE with the first ten genetic PCs, sex, genotyping batch and birth year as covariates and 240,000 pruned genetic markers for GRM estimation. Patients and control subjects in FinnGen provided informed consent for biobank research, based on the Finnish Biobank Act. Alternatively, older research cohorts, collected prior the start of FinnGen (in August 2017), were collected based on study-specific consents and later transferred to the Finnish biobanks after approval by Fimea, the National Supervisory Authority for Welfare and Health. Recruitment protocols followed the biobank protocols approved by Fimea. The Coordinating Ethics Committee of the Hospital District of Helsinki and Uusimaa (HUS) approved the FinnGen study protocol Nr HUS/990/2017.

The FinnGen study is approved by Finnish Institute for Health and Welfare (THL), approval number THL/2031/6.02.00/2017, amendments THL/1101/5.05.00/2017, THL/341/6.02.00/2018, THL/2222/6.02.00/2018, THL/283/6.02.00/2019, THL/1721/5.05.00/2019, Digital and population data service agency VRK43431/2017-3, VRK/6909/2018-3, VRK/4415/2019-3 the Social Insurance Institution (KELA) KELA 58/522/2017, KELA 131/522/2018, KELA 70/522/2019, KELA 98/522/2019, and Statistics Finland TK-53-1041-17. The Biobank Access Decisions for FinnGen samples and data utilized in FinnGen Data Freeze 5 include: THL Biobank BB2017_55, BB2017_111, BB2018_19, BB_2018_34, BB_2018_67, BB2018_71, BB2019_7, BB2019_8, BB2019_26, Finnish Red Cross Blood Service Biobank 7.12.2017, Helsinki Biobank HUS/359/2017, Auria Biobank AB17-5154, Biobank Borealis of Northern Finland_2017_1013, Biobank of Eastern Finland 1186/2018, Finnish Clinical Biobank Tampere MH0004, Central Finland Biobank 1-2017, and Terveystalo Biobank STB 2018001.

#### Genome build

The genomic coordinates reported in this paper were based on NCBI Build 37/UCSC hg19.

