## Supplementary tables and figures for "An efficient and accurate frailty model approach for genome-wide survival association analysis controlling for population structure and relatedness in large-scale biobanks"

*Supplementary Table 1: Projected computation time and memory usage for GATE and COXMEG-Score across different sample sizes. Benchmarking was performed for the GWAS of lifespan based on randomly subsampled data from UK Biobank White British ancestry subjects. Association tests were performed on 200,000 randomly selected markers with imputation INFO ≥ 0.3, with the filtering criteria of MAC ≥ 20. The computation times were projected for testing 46 million variants with INFO ≥ 0.3 and MAC ≥ 20. The reported run times are medians of five runs, each with randomly sampled subjects with different randomization seeds.*

| Sample Size (N) | GATE | | COXMEG-Score | |
|---|---|---|---|---|
| | Time (CPU-hr) | Memory (GB) | Time (CPU-hr) | Memory (GB) |
| 5000 | 27 | 0.36 | 192 | 1.80 |
| 10000 | 31 | 0.54 | 737 | 7.99 |
| 20000 | 34 | 0.74 | 3356 | 32.75 |
| 50000 | 59 | 1.44 | | |
| 100000 | 86 | 2.91 | | |
| 200000 | 147 | 5.55 | | |
| 408582 | 318 | 10.60 | | |

*Supplementary Table 2: Top genome-wide significant variants ($\alpha = 5\times10^{-8}$) in different loci based on GATE for four TTE phenotypes based on the UK Biobank data of subjects with White British ancestry and the Finngen study. For any variant with $p < 5\times10^{-8}$, we extend upstream and downstream by 1Mb, then merge the overlapping regions together to define the locus and report the variant that has the smallest p-value in each locus. Genomic coordinates are based on NCBI Build 37/UCSC hg19.*

| **Phenotype** | **Chr:Pos** | **rsID** | **Nearest Gene** | **Function** | **REF** | **ALT** | **UK Biobank** | | | | **FinnGen Study** | | | | **Meta-analysis UKBB + Finngen** | | **Previous case-control findings** |
|---|---|---|---|---|---|---|---|---|---|---|---|---|---|---|---|---|---|
| | | | | | | | **imputation info** | **AF** | **Hazard Ratio (95% CI)** | **p-value** | **imputation info** | **AF** | **Hazard Ratio (95% CI)** | **p-value** | **Hazard Ratio (95% CI)** | **p-value** | |
| Ischemic Heart Disease:<br><br>PheCode 411<br><br>UK Biobank N=407776, N events=36962, N censored=370814, censoring rate=90.9%<br><br>FinnGen Study N=218790, N events=30952, N censored=187838, censoring rate = 85.8% | 1:2984087 | rs2297829 | *PRDM16-DT* | ncRNA_exonic | C | A | 0.98 | 0.359 | 1.05(1.03-1.06) | 3.06E-08 | 0.98 | 0.464 | 1(0.98-1.02) | 8.64E-01 | 1.03(1.01-1.04) | 2.55E-05 | 1 |
| | 1:55505647 | rs11591147 | *PCSK9* | exonic | G | T | 1.00 | 0.018 | 0.79(0.74-0.84) | 9.64E-15 | 1.00 | 0.036 | 0.85(0.81-0.89) | 1.29E-10 | 0.83(0.8-0.86) | 5.83E-23 | 2,3 |
| | 1:56963627 | rs72664318 | *PLPP3* | intronic | A | G | 1.00 | 0.092 | 0.91(0.88-0.93) | 1.64E-12 | 1.00 | 0.109 | 0.93(0.9-0.95) | 1.24E-07 | 0.92(0.9-0.93) | 1.87E-18 | 2,4 |
| | 1:109817590 | rs12740374 | *CELSR2* | UTR3 | G | T | 1.00 | 0.221 | 0.9(0.88-0.92) | 1.54E-28 | 1.00 | 0.215 | 0.94(0.92-0.96) | 4.64E-09 | 0.92(0.9-0.93) | 2.57E-34 | 2,4 |
| | 1:222814442 | rs2133189 | *MIA3* | intronic | C | T | 1.00 | 0.714 | 1.07(1.05-1.09) | 5.80E-14 | 1.00 | 0.740 | 1.05(1.03-1.07) | 8.93E-06 | 1.06(1.05-1.07) | 7.95E-18 | 2,4 |
| | 2:19942473 | rs16986953 | *OSR1; LINC00954* | intergenic | G | A | 0.98 | 0.068 | 1.1(1.07-1.13) | 2.83E-09 | 1.00 | 0.074 | 1.06(1.02-1.1) | 7.77E-04 | 1.08(1.06-1.11) | 2.86E-11 | 2,4 |
| | 2:44073881 | rs6544713 | *ABCG8* | intronic | T | C | 1.00 | 0.676 | 0.95(0.94-0.97) | 3.26E-09 | 1.00 | 0.782 | 0.96(0.94-0.98) | 1.91E-04 | 0.95(0.94-0.97) | 3.22E-12 | 2,5 |
| | 2:85767735 | rs2028900 | *MAT2A* | intronic | C | T | 1.00 | 0.450 | 1.06(1.04-1.07) | 7.50E-12 | 1.00 | 0.401 | 1.04(1.03-1.06) | 2.99E-06 | 1.05(1.04-1.06) | 1.74E-16 | 2,4 |
| | 2:203865822 | rs145168080 | *CARF; NBEAL1* | intergenic | TGC | T | 0.99 | 0.125 | 1.09(1.06-1.11) | 9.34E-12 | NA | NA | NA | NA | NA | NA | 2,4 |
| | 3:136200780 | rs372408903 | *STAG1* | intronic | A | C | 0.99 | 0.129 | 0.93(0.91-0.96) | 9.68E-09 | NA | NA | NA | NA | NA | NA | 1 |
| | 3:138664512 | rs183837821 | *FOXL2* | exonic | G | A | 0.53 | 0.00010 | 132.95(23.19-762.32) | 4.09E-08 | NA | NA | NA | NA | NA | NA | 2,4 |
| | 4:81184341 | rs16998073 | *PRDM8; FGF5* | intergenic | A | T | 1.00 | 0.293 | 1.05(1.03-1.07) | 1.32E-08 | 1.00 | 0.312 | 1.02(1-1.04) | 2.71E-02 | 1.04(1.02-1.05) | 1.04E-08 | 6,7 |
| | 4:96117908 | rs376934295 | *UNC5C* | intronic | A | ATATATT | 0.97 | 0.787 | 0.95(0.93-0.96) | 1.47E-08 | NA | NA | NA | NA | NA | NA | 1 |
| | 4:110537251 | rs36121800 | *MCUB* | intronic | T | C | 0.95 | 0.018 | 1.2(1.13-1.28) | 3.52E-09 | 1.00 | 0.074 | 1.02(0.98-1.05) | 3.21E-01 | 1.06(1.03-1.09) | 0.000193 | - |
| | 4:148387701 | rs58721068 | *MIR548G; EDNRA* | intergenic | A | G | 0.99 | 0.143 | 1.07(1.04-1.09) | 6.96E-09 | 1.00 | 0.128 | 1.1(1.08-1.13) | 2.55E-13 | 1.08(1.06-1.1) | 6.58E-20 | 2 |
| | 4:156645513 | rs13139571 | *GUCY1A1* | intronic | C | A | 1.00 | 0.233 | 0.94(0.93-0.96) | 4.33E-10 | 1.00 | 0.233 | 0.94(0.92-0.96) | 6.14E-08 | 0.94(0.93-0.96) | 1.42E-16 | 2,4 |
| | 5:142516897 | rs246600 | *ARHGAP26* | intronic | C | T | 1.00 | 0.472 | 1.05(1.03-1.06) | 1.63E-08 | 1.00 | 0.480 | 1.03(1.01-1.05) | 4.29E-04 | 1.04(1.03-1.05) | 5.11E-11 | 8 |
| | 6:12903957 | rs9349379 | *PHACTR1* | intronic | A | G | 1.00 | 0.405 | 1.08(1.07-1.1) | 3.64E-23 | 1.00 | 0.451 | 1.09(1.07-1.11) | 5.40E-21 | 1.09(1.07-1.1) | 1.90E-42 | 2,4 |

| | | | | | | | | | | | | | | | | | |
|---|---|---|---|---|---|---|---|---|---|---|---|---|---|---|---|---|---|
| | 6:31585000 | rs2857597 | *AIF1* | downstream | T | A | 1.00 | 0.720 | 1.06(1.04-1.08) | 1.16E-10 | 1.00 | 0.799 | 1.03(1.01-1.05) | 9.44E-03 | 1.05(1.03-1.06) | 2.09E-11 | 2,4 |
| | 6:34604185 | rs3839632 | *ILRUN* | intronic | A | AT | 1.00 | 0.145 | 1.06(1.04-1.09) | 3.23E-08 | 1.00 | 0.207 | 1.01(0.98-1.03) | 6.34E-01 | 1.03(1.02-1.05) | 2.00E-05 | 1,2 |
| | 6:82437814 | rs117150895 | *TENT5A* | ncRNA_intronic | G | A | 1.00 | 0.022 | 0.84(0.79-0.88) | 1.51E-10 | 0.99 | 0.009 | 1.01(0.91-1.11) | 8.93E-01 | 0.88(0.84-0.92) | 3.16E-08 | - |
| | 6:134202690 | rs2327426 | *TARID* | ncRNA_intronic | T | C | 1.00 | 0.302 | 0.94(0.92-0.95) | 4.24E-14 | 1.00 | 0.246 | 0.96(0.94-0.98) | 9.14E-05 | 0.95(0.93-0.96) | 8.70E-17 | 2,4 |
| | 6:161005610 | rs55730499 | *LPA* | intronic | C | T | 1.00 | 0.081 | 1.29(1.25-1.33) | 1.77E-64 | 0.99 | 0.046 | 1.27(1.21-1.32) | 5.97E-26 | 1.28(1.25-1.31) | 1.48E-88 | 2,4 |
| | 7:19049388 | rs2107595 | *HDAC9; TWIST1* | intergenic | G | A | 0.99 | 0.152 | 1.07(1.05-1.09) | 1.67E-09 | 0.99 | 0.194 | 1.05(1.03-1.08) | 5.30E-06 | 1.06(1.05-1.08) | 6.50E-14 | 2,4 |
| | 7:150690176 | rs3918226 | *NOS3* | intronic | C | T | 0.97 | 0.081 | 1.11(1.07-1.14) | 1.16E-11 | 1.00 | 0.070 | 1.12(1.09-1.16) | 9.54E-11 | 1.11(1.09-1.14) | 8.60E-21 | 2,9 |
| | 8:4199406 | rs142333052 | *CSMD1* | intronic | C | G | 0.94 | 0.003 | 1.48(1.28-1.7) | 4.67E-08 | 0.99 | 0.021 | 1.03(0.96-1.09) | 4.23E-01 | 1.09(1.03-1.15) | 0.00321 | - |
| | 8:19855858 | rs200646600 | *LPL; SLC18A1* | intergenic | ATTTTT | A | 0.92 | 0.232 | 0.94(0.92-0.96) | 4.90E-10 | NA | NA | NA | NA | NA | NA | 2,4 |
| | 8:126504383 | rs10555326 | *TRIB1; LINC00861* | ncRNA_intronic | CCACCAT | C | 0.86 | 0.598 | 0.95(0.94-0.97) | 1.21E-08 | NA | NA | NA | NA | NA | NA | 2 |
| | 8:136004195 | rs544791777 | *LOC101927845; LINC01591* | intergenic | A | G | 0.61 | 0.00005 | 273.14(37-2016.66) | 3.95E-08 | NA | NA | NA | NA | NA | NA | - |
| | 9:22093299 | rs4007642 | *CDKN2B-AS1* | ncRNA_intronic | A | T | 0.99 | 0.484 | 1.17(1.15-1.19) | 6.42E-86 | NA | NA | NA | NA | NA | NA | 2,4 |
| | 10:30317073 | rs9337951 | *JCAD* | exonic | G | A | 0.95 | 0.345 | 1.05(1.04-1.07) | 1.24E-09 | 0.97 | 0.351 | 1.05(1.03-1.07) | 2.36E-06 | 1.05(1.04-1.06) | 1.63E-14 | 2,4 |
| | 10:44681231 | rs10626340 | *LINC00841; C10orf142* | intergenic | G | GCC | 1.00 | 0.115 | 0.93(0.9-0.95) | 4.21E-10 | NA | NA | NA | NA | NA | NA | 2,4 |
| | 11:9766932 | rs378825 | *SWAP70* | intronic | A | G | 1.00 | 0.573 | 1.04(1.03-1.06) | 4.92E-08 | 0.99 | 0.670 | 1.02(1-1.04) | 4.51E-02 | 1.03(1.02-1.05) | 4.19E-08 | 2,4 |
| | 11:24304661 | rs183283250 | *MIR8054; LUZP2* | intergenic | A | G | 0.36 | 0.00004 | 7.28E+06(2.67 E+04-1.98E+09) | 3.53E-08 | NA | NA | NA | NA | NA | NA | - |
| | 11:100593538 | rs633185 | *ARHGAP42* | intronic | G | C | 0.99 | 0.715 | 1.06(1.04-1.07) | 1.24E-09 | 1.00 | 0.711 | 1.02(1-1.04) | 1.32E-01 | 1.04(1.02-1.05) | 2.77E-08 | 1 |
| | 11:103673294 | rs2839812 | *DYNC2H1; MIR4693* | ncRNA_intronic | T | A | 1.00 | 0.721 | 0.94(0.92-0.95) | 6.40E-13 | 1.00 | 0.785 | 0.96(0.94-0.98) | 8.26E-05 | 0.95(0.93-0.96) | 6.85E-16 | 2,4 |
| | 11:107091429 | rs142065594 | *GUCY1A2, CWF19L2* | intergenic | T | TTG | 0.99 | 0.490 | 1.04(1.03-1.06) | 3.51E-08 | 1.00 | 0.545 | 1.01(0.99-1.03) | 2.44E-01 | 1.03(1.02-1.04) | 8.50E-07 | - |
| | 11:120237723 | rs368713737 | *ARHGEF12* | intronic | TA | T | 0.97 | 0.231 | 1.06(1.04-1.08) | 2.69E-10 | NA | NA | NA | NA | NA | NA | - |
| | 12:20168590 | rs796106347 | *LINC02398* | ncRNA_intronic | AT | A | 0.97 | 0.133 | 0.94(0.92-0.96) | 4.58E-08 | NA | NA | NA | NA | NA | NA | 7 |
| | 12:54513915 | rs11170820 | *FLJ12825* | ncRNA_exonic | C | G | 0.96 | 0.058 | 1.1(1.07-1.14) | 1.04E-08 | 1.00 | 0.060 | 1.1(1.06-1.14) | 1.08E-06 | 1.1(1.07-1.13) | 5.61E-14 | 6 |
| | 12:111904371 | rs4766578 | *ATXN2* | intronic | T | A | 1.00 | 0.505 | 0.93(0.92-0.95) | 1.76E-18 | 1.00 | 0.569 | 0.96(0.94-0.98) | 2.60E-06 | 0.94(0.93-0.95) | 2.75E-22 | 2,4 |
| | 12:121416650 | rs1169288 | *HNF1A* | exonic | A | C | 0.99 | 0.313 | 1.06(1.04-1.07) | 2.79E-10 | 1.00 | 0.374 | 1.02(1-1.04) | 3.57E-02 | 1.04(1.03-1.05) | 1.20E-09 | 6,7 |
| | 12:125303254 | rs112403212 | *SCARB1* | intronic | C | T | 0.98 | 0.139 | 1.07(1.04-1.09) | 2.87E-08 | NA | NA | NA | NA | NA | NA | 2,8 |
| | 13:110837553 | rs638634 | *COL4A1* | intronic | C | T | 0.95 | 0.302 | 0.95(0.93-0.97) | 1.67E-08 | 0.98 | 0.381 | 1(0.98-1.02) | 7.54E-01 | 0.97(0.96-0.98) | 1.47E-05 | 2,4 |

| | | | | | | | | | | | | | | | | | |
|---|---|---|---|---|---|---|---|---|---|---|---|---|---|---|---|---|---|
| | 14:100124055 | rs12897285 | *HHIPL1* | intronic | T | C | 1.00 | 0.171 | 0.94(0.92-0.96) | 2.77E-08 | NA | NA | NA | NA | NA | NA | 2,9 |
| | 15:79065380 | rs7177201 | *ADAMTS7* | intronic | T | C | 0.97 | 0.754 | 0.94(0.92-0.95) | 1.60E-12 | 1.00 | 0.647 | 0.93(0.92-0.95) | 5.11E-13 | 0.93(0.92-0.95) | 5.45E-24 | 2,4 |
| | 15:91427612 | rs12906125 | *FES* | intronic | G | A | 0.99 | 0.326 | 1.07(1.05-1.09) | 3.34E-16 | 1.00 | 0.271 | 1.04(1.01-1.06) | 7.84E-04 | 1.06(1.04-1.07) | 3.48E-17 | 2,4 |
| | 16:83045790 | rs7500448 | *CDH13* | intronic | A | G | 0.98 | 0.254 | 0.95(0.93-0.96) | 2.71E-09 | 0.99 | 0.205 | 0.96(0.93-0.98) | 4.54E-05 | 0.95(0.94-0.96) | 6.57E-13 | 6,7 |
| | 17:2038180 | rs1324063431 | *SMG6* | intronic | AT | A | 0.98 | 0.694 | 1.05(1.03-1.07) | 1.61E-08 | NA | NA | NA | NA | NA | NA | 1,2 |
| | 17:47488823 | rs55714120 | *PHB* | ncRNA_intronic | G | T | 0.99 | 0.338 | 1.06(1.04-1.08) | 8.18E-12 | 0.99 | 0.378 | 1.01(0.99-1.02) | 5.54E-01 | 1.03(1.02-1.05) | 4.16E-08 | 2,4 |
| | 19:11190292 | rs112032422 | *SMARCA4* | downstream | T | C | 1.00 | 0.118 | 0.91(0.88-0.93) | 1.99E-15 | 1.00 | 0.102 | 0.9(0.88-0.93) | 1.96E-11 | 0.91(0.89-0.92) | 2.60E-25 | 2,4 |
| | 19:18575193 | rs78030362 | *ELL* | intronic | A | G | 0.99 | 0.074 | 1.09(1.05-1.12) | 4.56E-08 | NA | NA | NA | NA | NA | NA | 1 |
| | 19:45412079 | rs7412 | *APOE* | exonic | C | T | 1.00 | 0.081 | 0.88(0.85-0.9) | 7.71E-19 | 1.00 | 0.054 | 0.84(0.81-0.88) | 3.61E-17 | 0.87(0.85-0.89) | 8.21E-34 | 2,4 |
| | 21:27355207 | rs150021565 | *APP* | intronic | A | G | 0.57 | 0.00021 | 21.12(7.17-62.18) | 3.02E-08 | NA | NA | NA | NA | NA | NA | - |
| | 21:35593827 | rs28451064 | *LINC00310; KCNE2* | ncRNA_intronic | G | A | 0.96 | 0.132 | 1.11(1.08-1.14) | 5.66E-18 | 0.99 | 0.153 | 1.1(1.07-1.13) | 3.17E-14 | 1.11(1.09-1.12) | 1.44E-30 | 2,4 |
| Female Breast Cancer: PheCode 174.1 UK Biobank N=208160, N events=15396, N censored=192764, censoring rate = 92.6% FinnGen Study N=123279, N events=8401, N censored=114878, censoring rate = 93.2% | 1:121280613 | rs11249433 | *EMBP1* | intronic | A | G | 1.00 | 0.417 | 1.08(1.06-1.11) | 6.32E-11 | 1.00 | 0.344 | 1.07(1.03-1.1) | 1.76E-04 | 1.08(1.06-1.1) | 6.05E-14 | 10 |
| | 2:121245613 | rs12616849 | *LINC01101; GLI2* | intergenic | G | C | 0.99 | 0.903 | 1.12(1.08-1.17) | 7.14E-09 | 0.99 | 0.935 | 1.14(1.07-1.21) | 1.20E-04 | 1.13(1.09-1.17) | 3.85E-12 | 11 |
| | 2:213537990 | rs3044147 | *ERBB4; LINC01878* | intergenic | TACCC | T | 0.99 | 0.365 | 1.07(1.05-1.1) | 7.32E-09 | 1.00 | 0.440 | 1.04(1.01-1.07) | 2.12E-02 | 1.06(1.04-1.08) | 1.86E-09 | 12 |
| | 2:217920769 | rs4442975 | *LOC101928278* | intronic | G | T | 0.99 | 0.512 | 0.88(0.86-0.90) | 2.63E-26 | 1.00 | 0.469 | 0.88(0.85-0.9) | 9.94E-16 | 0.88(0.86-0.90) | 2.21E-40 | 13 |
| | 3:27374101 | rs1352944 | *NEK10* | intronic | C | A | 1.00 | 0.476 | 0.9(0.88-0.92) | 1.21E-19 | 0.99 | 0.482 | 0.91(0.88-0.94) | 2.56E-08 | 0.9(0.89-0.92) | 2.53E-26 | 14 |
| | 4:175847144 | rs7654815 | *ADAM29* | intronic | A | T | 1.00 | 0.120 | 0.89(0.86-0.93) | 9.32E-10 | 1.00 | 0.130 | 0.91(0.87-0.95) | 7.99E-05 | 0.9(0.87-0.93) | 3.76E-13 | 11 |
| | 5:44706498 | rs10941679 | *LINC02224; MRPS30-DT* | intergenic | A | G | 0.98 | 0.254 | 1.14(1.11-1.17) | 1.15E-21 | 0.99 | 0.264 | 1.13(1.09-1.17) | 7.50E-11 | 1.14(1.11-1.16) | 6.82E-31 | 15 |
| | 5:46396206 | rs8188071 | *HCN1; NONE* | intergenic | G | A | 0.97 | 0.556 | 1.11(1.07-1.14) | 1.99E-10 | 0.99 | 0.669 | 0.97(0.94-1.01) | 1.06E-01 | 1.04(1.02-1.07) | 0.000274 | 16 |
| | 5:56007339 | rs7709971 | *C5orf67; LOC105378979* | intergenic | G | A | 1.00 | 0.162 | 1.2(1.17-1.24) | 2.11E-29 | 1.00 | 0.133 | 1.16(1.1-1.21) | 1.78E-09 | 1.19(1.16-1.22) | 6.42E-37 | 17 |
| | 5:158227898 | rs12332693 | *EBF1* | intronic | G | A | 0.99 | 0.548 | 0.92(0.90-0.94) | 5.74E-12 | 1.00 | 0.592 | 0.94(0.91-0.97) | 3.87E-04 | 0.93(0.91-0.95) | 1.80E-14 | 11 |
| | 6:82355896 | rs12214288 | *BCKDHB; TENT5A* | intergenic | A | G | 0.98 | 0.213 | 1.08(1.05-1.12) | 3.21E-08 | 1.00 | 0.287 | 1.06(1.02-1.1) | 1.68E-03 | 1.07(1.05-1.1) | 3.53E-10 | 18 |
| | 6:151949806 | rs6913578 | *CCDC170; ESR1* | intergenic | A | C | 1.00 | 0.323 | 1.13(1.1-1.15) | 2.07E-20 | 1.00 | 0.195 | 1.12(1.08-1.17) | 3.42E-08 | 1.12(1.1-1.15) | 4.42E-27 | 19 |
| | 8:36859186 | rs12681990 | *KCNU1; LINC01605* | intergenic | T | C | 0.99 | 0.158 | 0.9(0.87-0.93) | 5.15E-11 | 1.00 | 0.141 | 0.87(0.83-0.91) | 7.33E-09 | 0.89(0.87-0.91) | 3.68E-18 | 20 |
| | 8:98847428 | rs549309990 | *LAPTM4B* | intronic | C | T | 0.70 | 0.00033 | 15.18(5.72-40.29) | 4.76E-08 | NA | NA | NA | NA | NA | NA | 21 |
| | 8:128356670 | rs17465317 | *CASC21,CASC8* | ncRNA_intronic | C | T | 1.00 | 0.545 | 1.10(1.08-1.13) | 1.40E-16 | NA | NA | NA | NA | NA | NA | 22 |

| | | | | | | | | | | | | | | | | | |
|---|---|---|---|---|---|---|---|---|---|---|---|---|---|---|---|---|---|
| | 9:110893720 | rs34138847 | *KLF4; ACTL7B* | intergenic | GTATT | G | 0.98 | 0.624 | 1.10(1.07-1.12) | 1.60E-13 | 1.00 | 0.662 | 1.12(1.08-1.16) | 2.93E-11 | 1.10(1.08-1.13) | 5.77E-23 | 23 |
| | 10:21799726 | rs12256551 | *MIR1915HG; SKIDA1* | intergenic | A | C | 0.99 | 0.353 | 1.08(1.05-1.1) | 3.59E-09 | NA | NA | NA | NA | NA | NA | 16 |
| | 10:64288130 | rs10995194 | *ZNF365* | ncRNA_intronic | G | C | 0.99 | 0.144 | 0.9(0.87-0.93) | 2.16E-09 | 1.00 | 0.202 | 0.9(0.87-0.94) | 1.14E-06 | 0.9(0.88-0.93) | 1.23E-14 | 24 |
| | 10:80887430 | rs11269768 | *ZMIZ1* | intronic | GGCCCCTGCTCA | G | 0.99 | 0.158 | 1.11(1.08-1.15) | 1.26E-10 | 1.00 | 0.140 | 1.09(1.04-1.14) | 1.85E-04 | 1.1(1.08-1.13) | 1.22E-13 | 25 |
| | 10:123348662 | rs2912774 | *FGFR2* | intronic | T | G | 1.00 | 0.605 | 0.79(0.77-0.81) | 1.94E-83 | 1.00 | 0.564 | 0.82(0.8-0.85) | 1.53E-31 | 0.80(0.79-0.82) | 3.19E-112 | 26 |
| | 11:1898664 | rs1973765 | *LSP1* | intronic | T | C | 0.99 | 0.387 | 0.93(0.9-0.95) | 4.89E-10 | 0.99 | 0.414 | 0.91(0.88-0.94) | 8.24E-08 | 0.92(0.9-0.94) | 2.65E-16 | 22 |
| | 11:69331642 | rs554219 | *LINC01488; CCND1* | intergenic | C | G | 0.99 | 0.120 | 1.26(1.22-1.31) | 1.96E-36 | 1.00 | 0.148 | 1.25(1.2-1.31) | 5.93E-22 | 1.26(1.23-1.3) | 1.20E-56 | 27 |
| | 11:129474008 | rs1491531488 | *RPS27P20; LINC01395* | intergenic | TAA | T | 0.99 | 0.545 | 1.07(1.05-1.10) | 2.13E-09 | NA | NA | NA | NA | NA | NA | 11 |
| | 12:25518873 | rs10842542 | *KRAS; LMNTD1* | intergenic | T | G | 1.00 | 0.452 | 0.94(0.91-0.96) | 2.26E-08 | 0.99 | 0.350 | 0.99(0.96-1.02) | 5.63E-01 | 0.95(0.93-0.97) | 8.33E-07 | - |
| | 12:28151609 | rs812020 | *PTHLH; LOC729291* | intergenic | A | C | 0.97 | 0.260 | 0.92(0.89-0.94) | 6.55E-10 | 1.00 | 0.247 | 0.91(0.88-0.95) | 2.44E-06 | 0.92(0.9-0.94) | 8.10E-15 | 28 |
| | 12:96026737 | rs61938093 | *USP44; PGAM1P5* | intergenic | C | T | 0.99 | 0.295 | 0.9(0.87-0.92) | 1.01E-16 | 0.99 | 0.347 | 0.89(0.86-0.92) | 4.14E-11 | 0.9(0.88-0.91) | 2.87E-26 | 11 |
| | 12:115834946 | rs2133317 | *TBX3; MED13L* | intergenic | C | G | 1.00 | 0.385 | 0.92(0.9-0.95) | 1.11E-10 | 1.00 | 0.382 | 0.94(0.91-0.97) | 4.89E-04 | 0.93(0.91-0.95) | 3.66E-13 | 29 |
| | 14:68978984 | rs35378451 | *RAD51B* | intronic | CA | C | 0.97 | 0.284 | 0.91(0.89-0.93) | 1.93E-12 | 0.99 | 0.245 | 0.91(0.88-0.95) | 3.10E-06 | 0.91(0.89-0.93) | 3.11E-17 | 10 |
| | 16:52599188 | rs4784227 | *CASC16* | ncRNA_intronic | C | T | 1.00 | 0.239 | 1.26(1.23-1.3) | 6.38E-60 | 1.00 | 0.251 | 1.25(1.21-1.3) | 9.81E-32 | 1.26(1.23-1.29) | 7.85E-90 | 30 |
| | 16:53824226 | rs62033406 | *FTO* | intronic | A | G | 1.00 | 0.407 | 0.93(0.91-0.95) | 5.31E-10 | 1.00 | 0.441 | 0.94(0.91-0.97) | 3.07E-04 | 0.93(0.92-0.95) | 9.09E-13 | 11 |
| | 16:80651180 | rs879843 | *CDYL2* | intronic | A | G | 1.00 | 0.219 | 1.09(1.06-1.12) | 5.46E-09 | 1.00 | 0.270 | 1.07(1.03-1.11) | 5.71E-04 | 1.08(1.06-1.1) | 1.80E-11 | 11 |
| | 17:44321478 | rs2732714 | *KANSL1; ARL17B* | intergenic | A | G | 0.92 | 0.196 | 0.92(0.89-0.94) | 1.96E-08 | NA | NA | NA | NA | NA | NA | 16 |
| | 18:24624030 | rs138736090 | *CHST9* | intronic | C | CT | 0.94 | 0.149 | 0.9(0.87-0.93) | 2.14E-09 | 0.98 | 0.062 | 1.01(0.94-1.07) | 8.79E-01 | 0.92(0.9-0.95) | 1.39E-07 | 11 |
| | 21:16563640 | rs2823129 | *NRIP1* | intergenic | C | T | 0.99 | 0.324 | 0.93(0.91-0.95) | 9.56E-09 | 1.00 | 0.304 | 0.91(0.88-0.95) | 6.34E-07 | 0.92(0.91-0.94) | 4.03E-14 | 29 |
| | 22:28314612 | rs62235635 | *PITPNB* | intronic | G | A | 0.80 | 0.008 | 1.75(1.5-2.05) | 1.91E-12 | 0.97 | 0.017 | 1.65(1.45-1.88) | 1.09E-13 | 1.69(1.53-1.87) | 1.63E-24 | 11 |
| | 22:29450193 | rs62236881 | *ZNRF3* | UTR3 | G | A | 0.78 | 0.003 | 4.71(3.49-6.36) | 4.39E-24 | 0.99 | 0.008 | 3(2.45-3.68) | 6.13E-27 | 3.46(2.92-4.09) | 4.89E-48 | 11 |
| | 22:40935623 | rs147759720 | *MRTFA* | intronic | C | T | 0.89 | 0.079 | 1.2(1.15-1.26) | 1.65E-14 | NA | NA | NA | NA | NA | NA | 11 |
| Glaucoma: | 1:97354095 | rs568063121 | *PTBP2; DPYD* | intergenic | T | C | 0.67 | 0.00024 | 1012.32(87.36-11731.12) | 2.83E-08 | NA | NA | NA | NA | NA | NA | - |
| PheCode 365 | 1:165708633 | rs5778472 | *TMCO1* | intronic | A | AT | 0.99 | 0.876 | 0.75(0.71-0.8) | 6.71E-22 | NA | NA | NA | NA | NA | NA | 31 |
| UK Biobank N=398971, N events=6046, | 1:171605478 | rs74315329 | *MYOC* | exonic | G | A | 0.94 | 0.001 | 28.79(14.47-57.28) | 8.84E-22 | 1.00 | 0.003 | 4.62(3.26-6.53) | 5.69E-18 | 6.69(4.91-9.11) | 2.51E-33 | 32 |
| N censored=392925, censoring rate = 98.5% | 1:173008989 | rs779440307 | *AIMP1P2; TNFSF18* | intergenic | C | T | 0.37 | 0.000 | 221.41(31.99-1532.29) | 4.55E-08 | NA | NA | NA | NA | NA | NA | - |
| FinnGen Study | 3:85134557 | rs9309969 | *CADM2* | intronic | T | G | 0.99 | 0.594 | 0.87(0.84-0.90) | 1.48E-13 | 1.00 | 0.475 | 0.95(0.92-0.98) | 8.65E-04 | 0.91(0.89-0.94) | 2.25E-13 | 33 |

| | | | | | | | | | | | | | | | | | |
|---|---|---|---|---|---|---|---|---|---|---|---|---|---|---|---|---|---|
| N=218790,<br>N events=8591,<br>N censored=210199,<br>censoring rate = 96.1% | 3:186128816 | rs56233426 | *DGKG; LINC02020* | intergenic | A | G | 0.96 | 0.537 | 0.88(0.85-0.92) | 8.46E-11 | 0.98 | 0.515 | 0.94(0.91-0.97) | 4.56E-05 | 0.91(0.89-0.94) | 2.28E-13 | 34,35 |
| | 4:7918390 | rs16841407 | *AFAP1* | intronic | G | A | 0.99 | 0.317 | 1.17(1.13-1.22) | 2.23E-15 | 0.99 | 0.347 | 1.12(1.08-1.16) | 1.06E-10 | 1.14(1.11-1.17) | 8.52E-24 | 36 |
| | 4:169237230 | rs755949313 | *DDX60* | intronic | C | T | 0.58 | 0.00015 | 5014.05(240.33-104610.58) | 3.74E-08 | NA | NA | NA | NA | NA | NA | - |
| | 7:11677840 | rs2526099 | *THSD7A* | intronic | A | G | 0.97 | 0.473 | 1.11(1.07-1.16) | 9.58E-09 | 0.99 | 0.414 | 1.04(1.01-1.08) | 9.93E-03 | 1.07(1.05-1.1) | 1.05E-08 | 34,35 |
| | 7:116153329 | rs10281661 | *CAV2; CAV1* | intergenic | A | G | 1.00 | 0.273 | 1.12(1.08-1.17) | 4.09E-08 | 1.00 | 0.149 | 1.09(1.04-1.14) | 1.86E-04 | 1.11(1.07-1.14) | 5.02E-11 | 33 |
| | 7:146348027 | rs540694424 | *CNTNAP2* | intronic | G | C | 0.69 | 0.00004 | 7.86E+15(1.93E+10-3.20E+21) | 2.71E-08 | NA | NA | NA | NA | NA | NA | - |
| | 9:22056295 | rs7853090 | *CDKN2B-AS1* | ncRNA_exonic | T | C | 1.00 | 0.568 | 1.19(1.14-1.23) | 1.06E-19 | 1.00 | 0.560 | 1.10(1.06-1.13) | 3.07E-08 | 1.13(1.11-1.16) | 3.79E-24 | 31 |
| | 9:107693201 | rs2437812 | *ABCA1; SLC44A1* | intergenic | A | C | 0.98 | 0.576 | 0.85(0.82-0.88) | 1.72E-18 | NA | NA | NA | NA | NA | NA | 36 |
| | 9:129373110 | rs12377624 | *MVB12B; LMX1B* | ncRNA_intronic | G | C | 1.00 | 0.372 | 0.89(0.86-0.93) | 4.29E-09 | 1.00 | 0.383 | 0.87(0.84-0.9) | 5.90E-16 | 0.88(0.86-0.9) | 2.32E-23 | 33 |
| | 11:47275064 | rs10838681 | *NR1H3* | intronic | G | A | 1.00 | 0.268 | 1.14(1.09-1.18) | 2.00E-09 | 1.00 | 0.354 | 1.08(1.04-1.12) | 9.38E-06 | 1.1(1.07-1.13) | 4.55E-13 | 33 |
| | 11:120316907 | rs59459827 | *ARHGEF12* | intronic | AAAT | A | 0.91 | 0.872 | 0.84(0.80-0.89) | 5.70E-09 | NA | NA | NA | NA | NA | NA | 36 |
| | 15:28365618 | rs12913832 | *HERC2* | intronic | A | G | 1.00 | 0.786 | 0.88(0.84-0.92) | 2.17E-08 | NA | NA | NA | NA | NA | NA | 37 |
| | 17:10031090 | rs12150284 | *GAS7* | intronic | C | T | 1.00 | 0.373 | 0.86(0.83-0.9) | 1.68E-14 | 1.00 | 0.374 | 0.88(0.85-0.91) | 4.43E-15 | 0.87(0.85-0.89) | 6.34E-28 | 33,38 |
| | 17:58934999 | rs9895116 | *BCAS3* | intronic | T | G | 1.00 | 0.146 | 1.16(1.11-1.23) | 1.15E-08 | 1.00 | 0.142 | 1.05(1-1.1) | 4.59E-02 | 1.1(1.06-1.13) | 1.41E-07 | 33 |
| Alzheimer's Disease:<br><br>PheCode 290.11<br><br>UK Biobank<br>N=342881,<br>N events=822,<br>N censored=342059,<br>censoring rate = 99.8%<br><br>FinnGen Study<br>N=207713,<br>N events=3899,<br>N censored=207324,<br>censoring rate = 93.2% | 10:37029593 | rs749680474 | *PCAT5; ANKRD30A* | intergenic | A | G | 0.54 | 0.00060 | 2.97E+05(3.75E+03-2.35E+07) | 1.38E-08 | NA | NA | NA | NA | NA | NA | - |
| | 18:76901532 | rs533100590 | *ATP9B* | intronic | T | C | 0.33 | 0.00005 | 4.33E+91(1.95E+59-9.60E+123) | 2.78E-08 | NA | NA | NA | NA | NA | NA | - |
| | 19:45411941 | rs429358 | *APOE* | exonic | T | C | 1.00 | 0.155 | 6.75(5.73-7.96) | 7.02E-115 | 1.00 | 0.182 | 3.74(3.48-4.02) | 6.13E-282 | 4.1(3.83-4.37) | 4.94E-324 | 39 |

*Supplementary Table 3: Top genome-wide significant variants ($\alpha = 5\times10^{-8}$) in different loci based on GATE for lifespan based on the FinnGen Study and the UK Biobank data. For any variant with $p < 5\times10^{-8}$, we extend upstream and downstream by 1Mb, then merge the overlapping regions together to define the locus and report the variant that has the smallest p-value in each locus. Genomic coordinates are based on NCBI Build 37/UCSC hg19.*

| Phenotype | Chr:Pos | rsID | Nearest Gene | Function | REF | ALT | UK Biobank | | | | Finngen Study | | | | Meta-analysis UKBB + FinnGen | |
|---|---|---|---|---|---|---|---|---|---|---|---|---|---|---|---|---|
| | | | | | | | Imputation INFO | AF | Hazard Ratio (95% CI) | p-value | Imputation INFO | AF | Hazard Ratio (95% CI) | p-value | Hazard Ratio (95% CI) | p-value |
| Lifespan<br><br>UK Biobank<br>N=406596,<br>N events=16875,<br>N censored=389721,<br>censoring rate = 95.8%<br><br>FinnGen study<br>N=218396,<br>N events=15152,<br>N censored=203244,<br>censoring rate = 93.1% | 19: 45424514 | rs157592* | *APOC1; APOC1P1* | intergenic | A | C | 0.95 | 0.187 | 1.08 (1.05, 1.12) | 1.87E-08 | | | | Not found in FinnGen | NA | NA |
| | 19:45411941 | rs429358* | *APOE* | missense | T | C | 1 | 0.156 | 1.07 (1.04, 1.10) | 1.92E-05 | 1 | 0.183 | 1.13 (1.10, 1.17) | 1.01E-14 | 1.1(1.07-1.12) | 4.04E-17 |

* rs157592 and rs429358 are in LD with $r^2 = 0.7$ in UK Biobank

*Supplementary Table 4: Empirical type I error rates of GATE and GATE with no SPA based on $9.4x10^8$ association tests in 100 simulated data sets with censoring rates 50%, 75% and 90%, respectively. Each data set contains 5,000 independent individuals and 500 families. Each family was simulated following the pedigree structure shown in Supplementary Figure 7. The variance component parameter $\tau$ = 0.1 and 0.25.*

| Variance component parameter $\tau$ | | Alpha | Censoring Rate | | |
|---|---|---|---|---|---|
| | | | 50% | 75% | 90% |
| 0.1 | SPA | 1.00E-06 | 1.21E-06 | 1.11E-06 | 1.03E-06 |
| | | 5.00E-08 | 7.41E-08 | 5.81E-08 | 5.22E-08 |
| | no SPA | 1.00E-06 | 1.27E-05 | 2.07E-05 | 5.12E-05 |
| | | 5.00E-08 | 4.79E-06 | 8.90E-06 | 2.81E-05 |
| 0.25 | SPA | 1.00E-06 | 1.21E-06 | 1.08E-06 | 9.72E-07 |
| | | 5.00E-08 | 6.12E-08 | 5.10E-08 | 5.53E-08 |
| | no SPA | 1.00E-06 | 1.20E-05 | 1.91E-05 | 4.67E-05 |
| | | 5.00E-08 | 4.38E-06 | 8.18E-06 | 2.50E-05 |

*Supplementary Table 5: Empirical type I error rates of GATE and GATE with no SPA based on 8.3x10$^{8}$ association tests in 100 simulated data sets with censoring rates 50%, 75% and 90%, respectively. Each data set contains 10,000 randomly selected individuals with white British ancestry from the UK Biobank. The variance component parameter τ = 0.25.*

| | | Censoring rate | | |
|---|---|---|---|---|
| | **Alpha** | **50%** | **75%** | **90%** |
| **SPA** | **1.00E-06** | 1.20E-06 | 1.09E-06 | 8.94E-07 |
| | **5.00E-08** | 7.09E-08 | 5.65E-08 | 3.85E-08 |
| **no SPA** | **1.00E-06** | 1.12E-05 | 1.90E-05 | 6.55E-05 |
| | **5.00E-08** | 4.35E-06 | 8.38E-06 | 3.99E-05 |

### Supplementary Figures

*Supplementary Figure 1: Comparing association p-values from GATE versus COXMEG based on 5 million genetic variants in simulation data sets. A. Scatter plots of association p-values from GATE versus COXMEG. B. Quantile-quantile plots stratified by MAF for GATE and COXMEG. For each censoring rate, 100 data sets were simulated, each has 10,000 samples (5,000 independent samples and 500 families, each with 10 family members as shown in Supplementary Figure 7). For each data set, 50,000 simulated genetic markers were tested. C. . Scatter plots of association p-values from GATE-noSPA versus COXMEG.*

A.

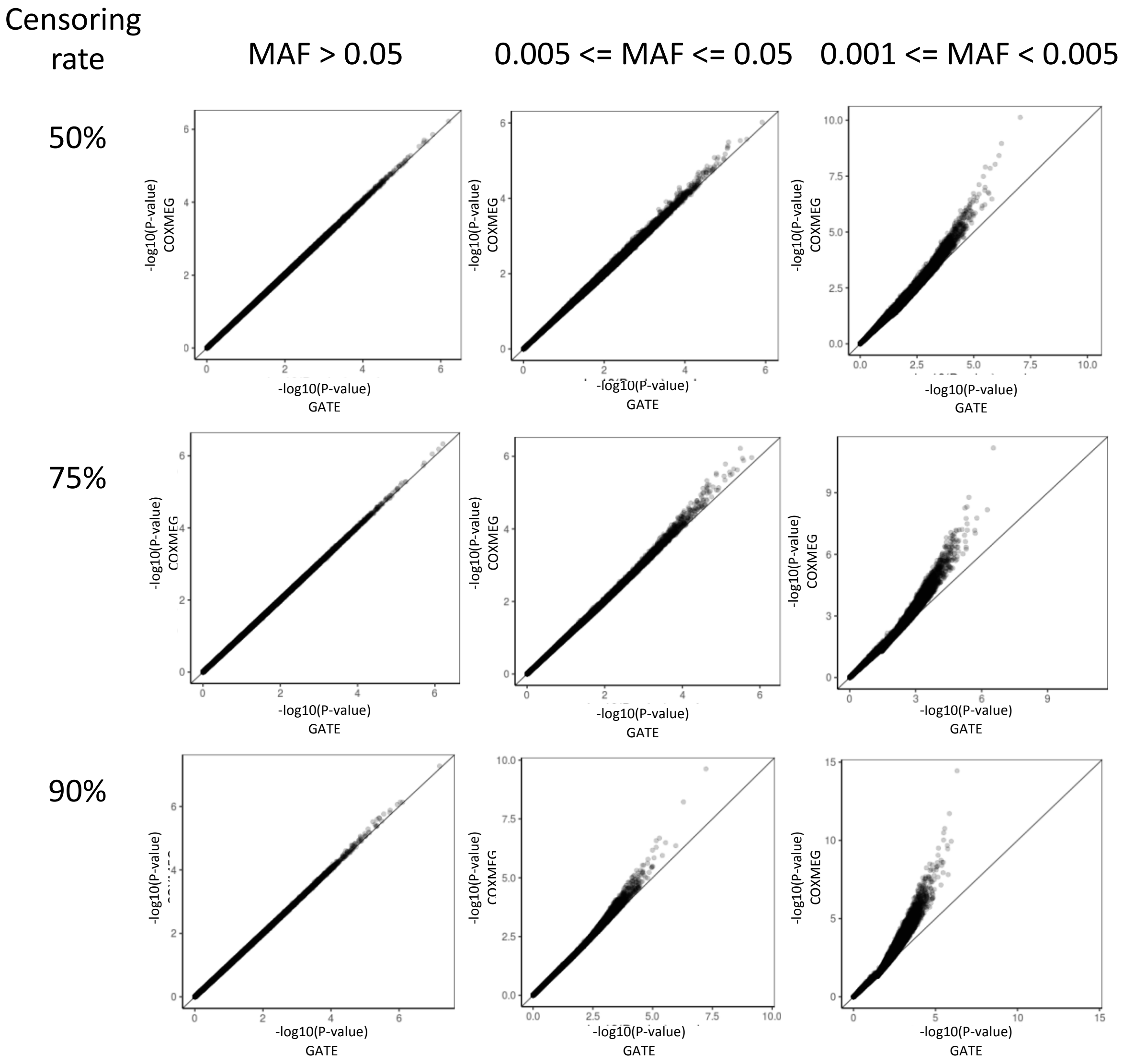

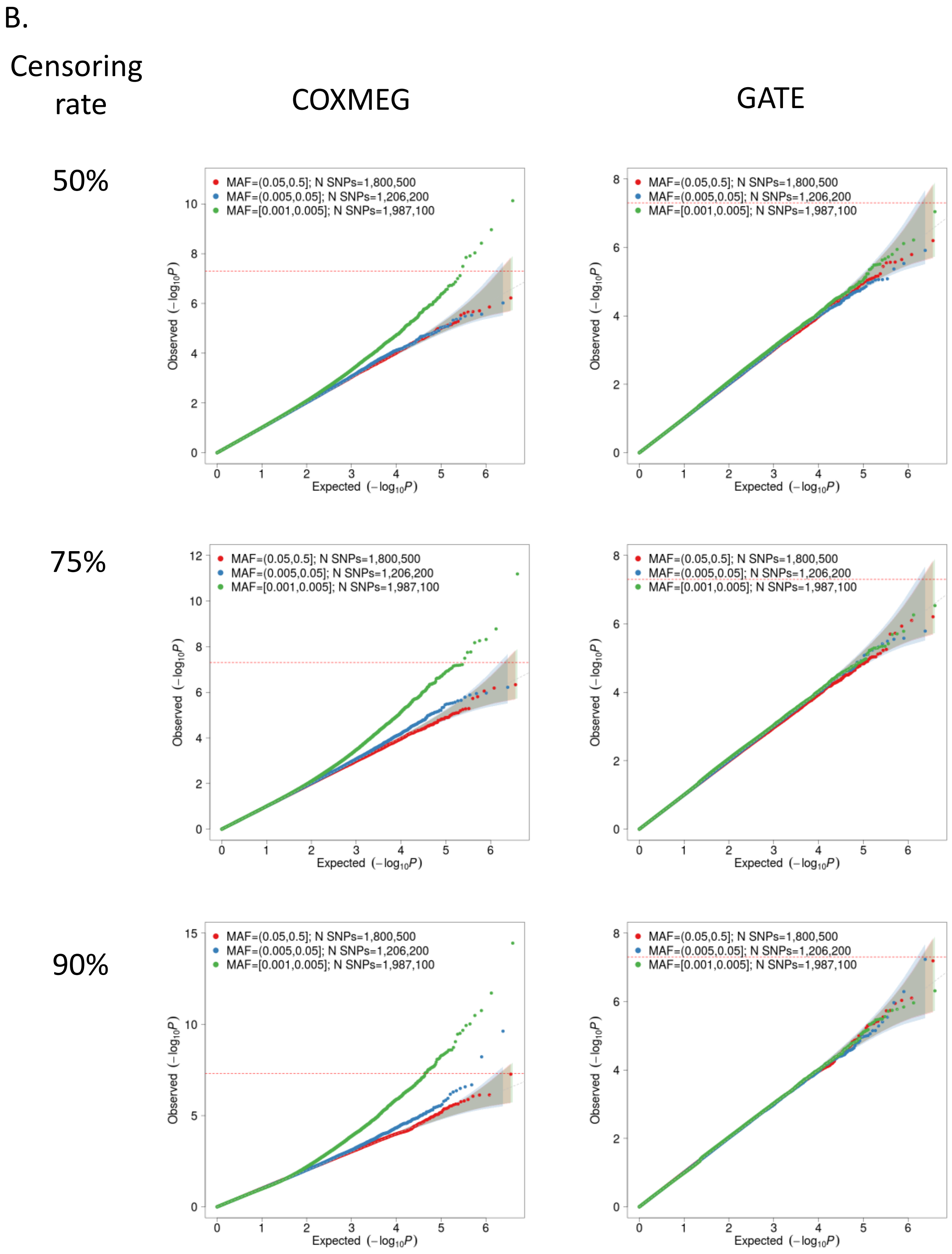

B.
Censoring rate
COXMEG
GATE
50%
75%
90%
MAF=(0.05,0.5]; N SNPs=1,800,500
MAF=(0.005,0.05]; N SNPs=1,206,200
MAF=[0.001,0.005]; N SNPs=1,987,100
Observed $(-\log_{10}P)$
Expected $(-\log_{10}P)$

C.

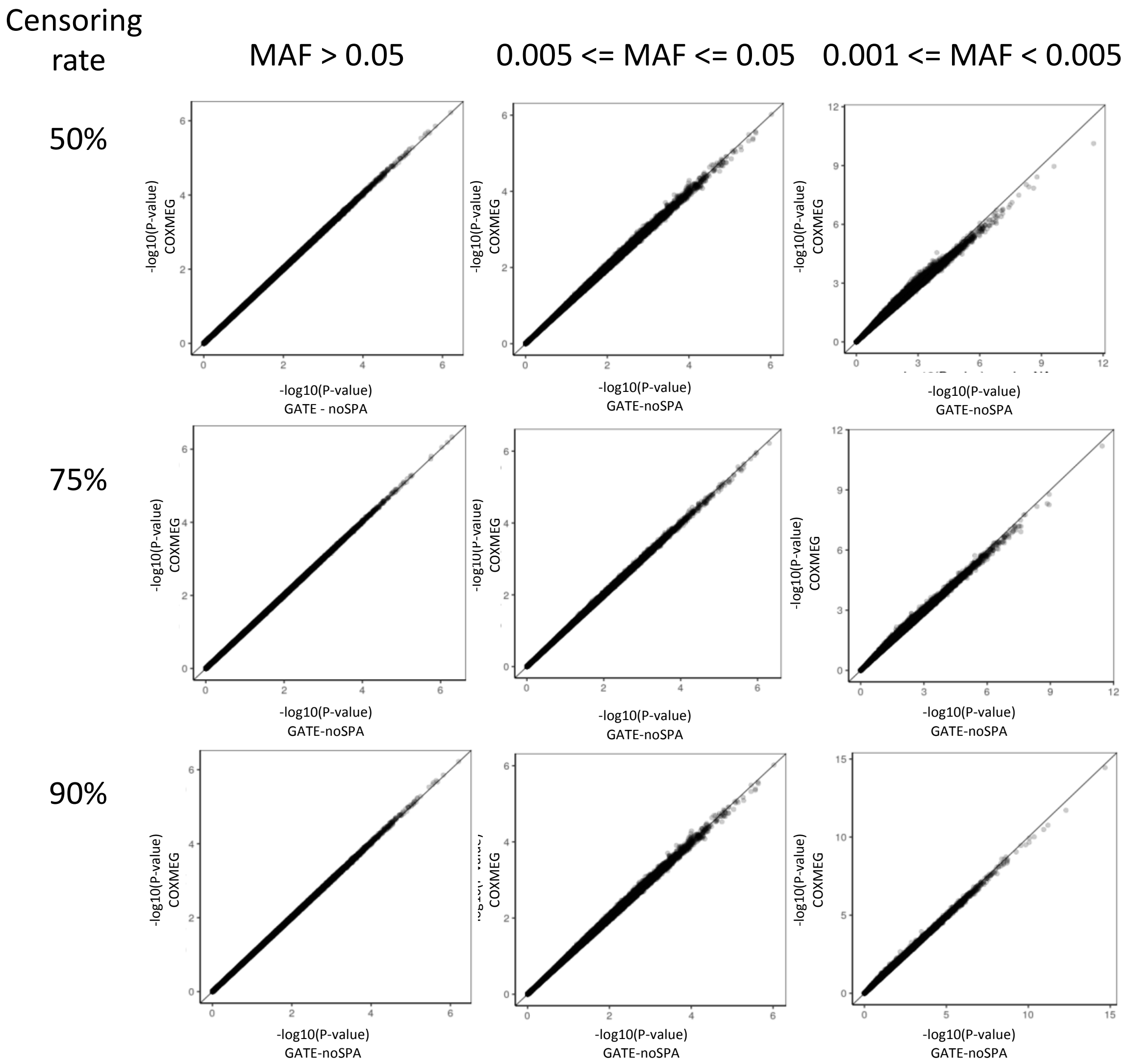

Censoring rate
MAF > 0.05
0.005 <= MAF <= 0.05
0.001 <= MAF < 0.005
50%
75%
90%
-log10(P-value) COXMEG
-log10(P-value) GATE - noSPA
-log10(P-value) GATE-noSPA

*Supplementary Figure 2: Histogram of censoring rates of 871 PheCodes in the UK Biobank subjects with White British ancestry. The PheCodes are constructed based on the ICD9 and ICD10 codes and the associated diagnostic dates. Detailed description of the PheCode construction is available in the ONLINE METHODS section.*

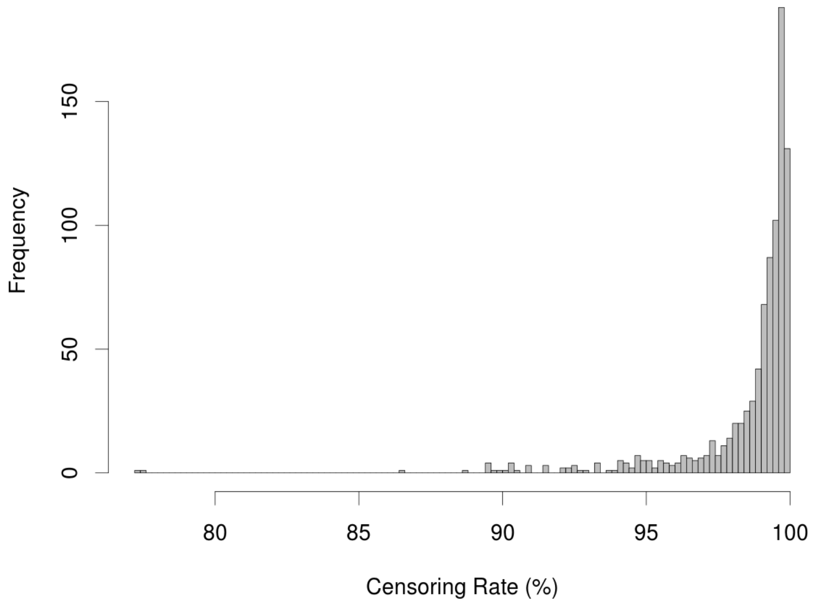

*Supplementary Figure 3: Disease-free probability over-time by genotypes for loci LPA and CELSR2 for ischemic heart disease, FGFR2 and CASC16 for female breast cancer, MYOC and TMCO1 for glaucoma, and APOE e4 variant for AD. The red, green and blue lines represent the disease-free probability for alternate allele counts zero, one and two, respectively.*

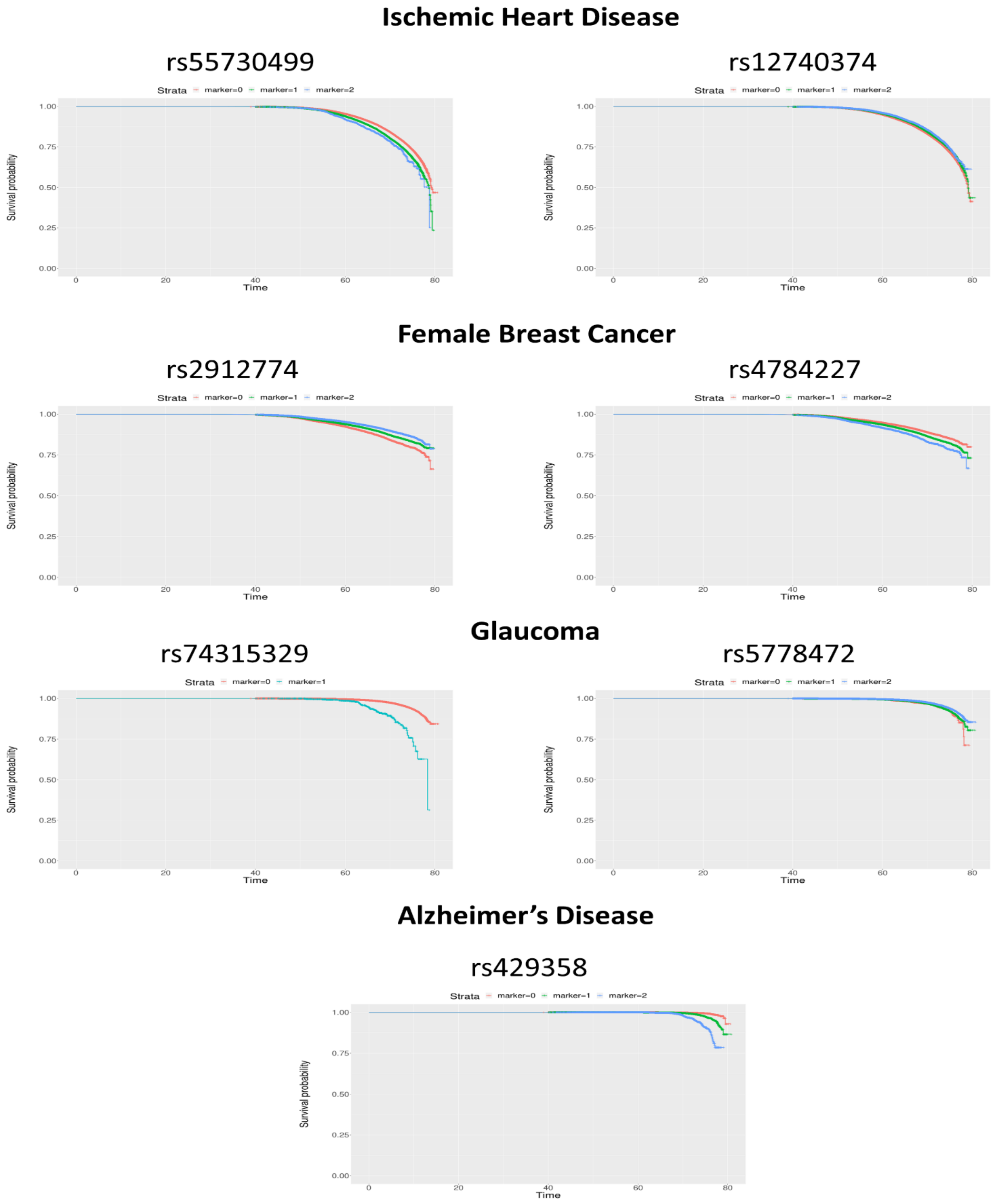

*Supplementary Figure 4: GWAS of lifespan in the FinnGen Study (N events=15152, N censored=203244): A. Overall survival curve for lifespan B. GATE QQ plot (left) and Manhattan plot (right).*

A.

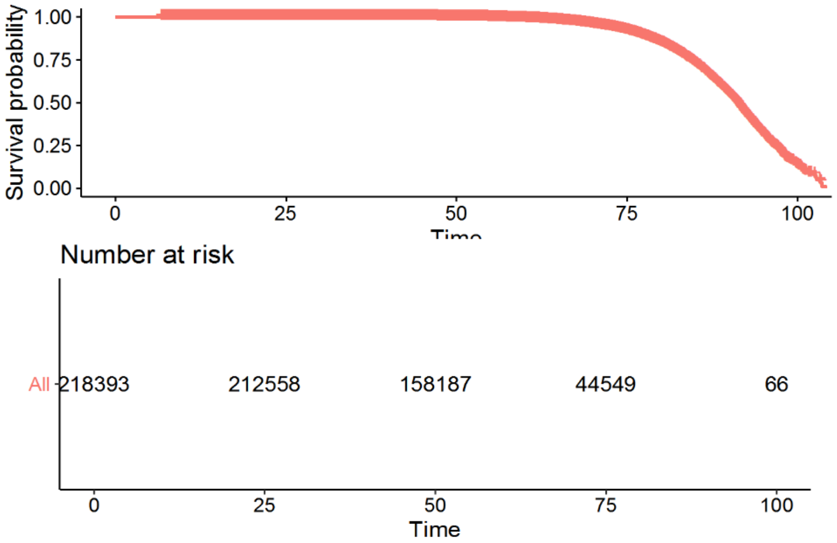


B. GATE

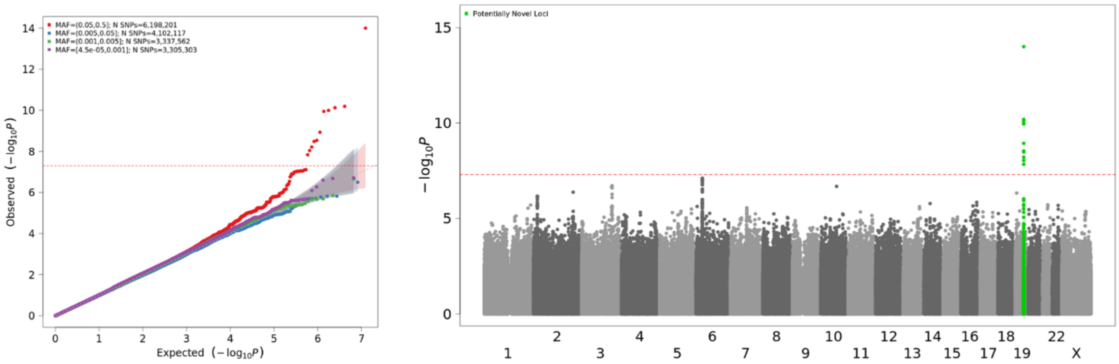

*Supplementary Figure 5: GWAS of lifespan in the UK Biobank (N events=16875, N censored=389721): A. Overall survival curve for lifespan B. GATE QQ plot (left) and Manhattan plot (right).*

A.

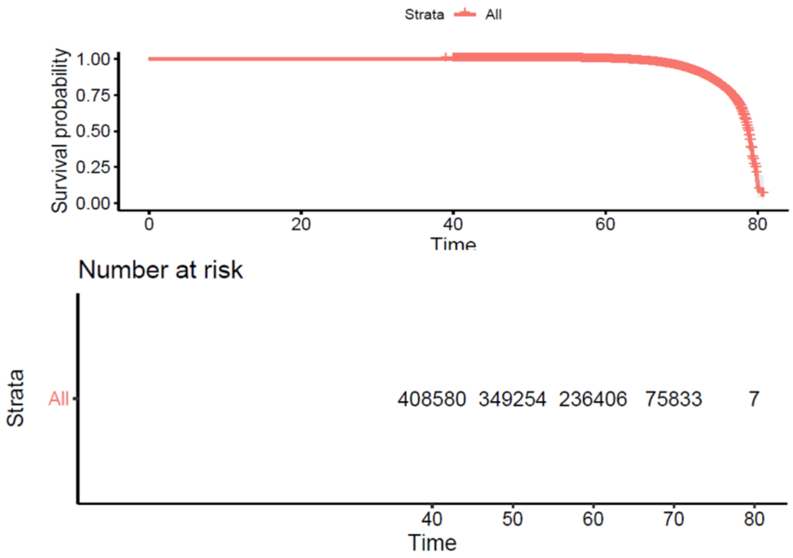


B. GATE

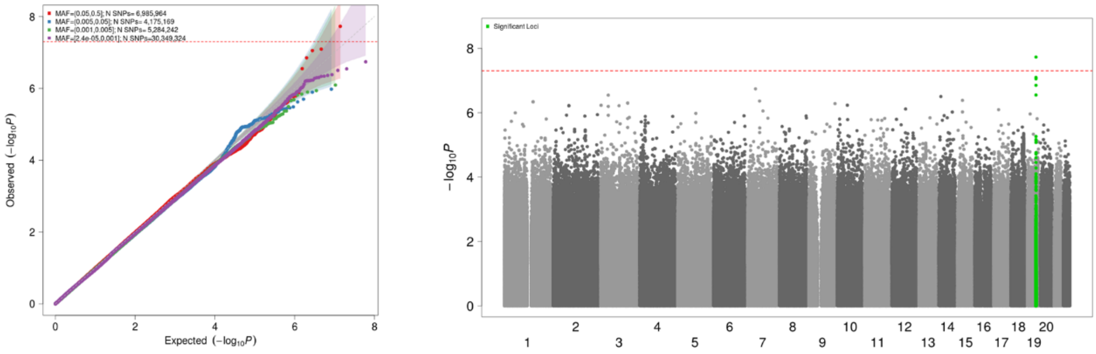

*Supplementary Figure 6: Meta-analysis Manhattan plot of the GWAS results for lifespan in the UK Biobank (N events=16875, N censored=389721) and FinnGen Study (N events=15152, N censored=203244).*

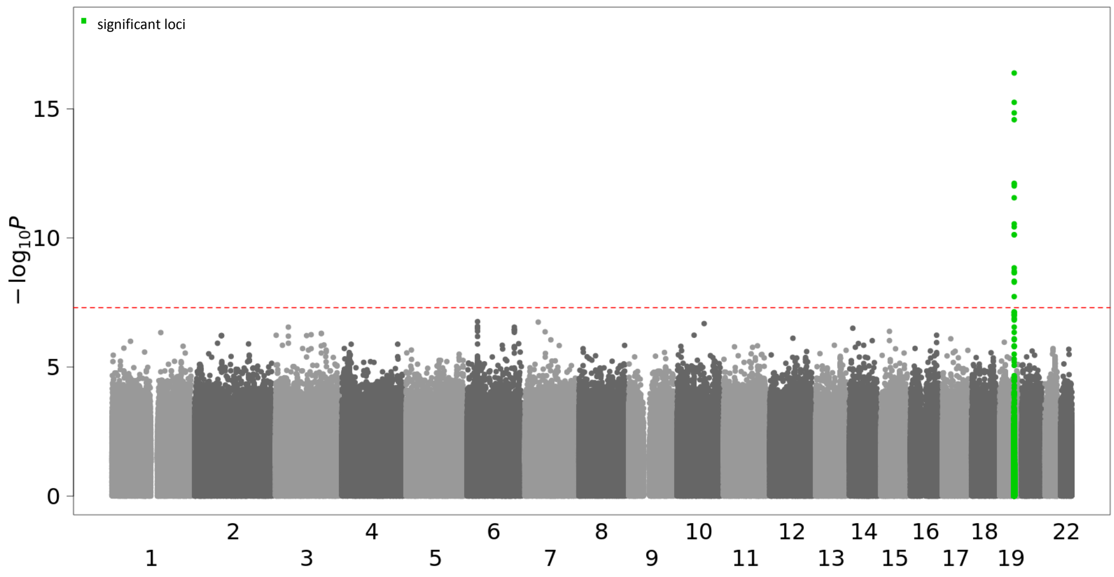


*Supplementary Figure 7: Pedigree of families, each with 10 members, in the simulation study.*

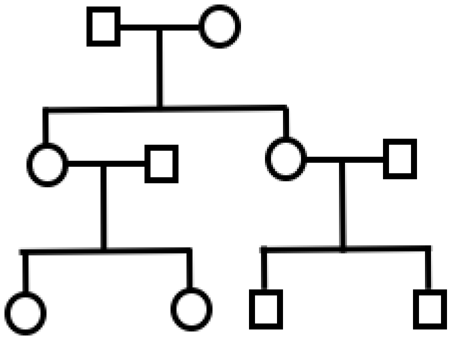

*Supplementary Figure 8: Empirical type I error rates and 95% confidence intervals (horizontal bars) for GATE and GATE with no SPA estimated in simulation studies were plotted for censoring rates 50%. 75%,*

*and 90%. A. based 9.4x10*$^{8}$ *association tests. For each censoring rate, 100 data sets were simulated. Each data set contains 5,000 independent individuals and 500 families. Each family was simulated following the pedigree structure shown in Supplementary Figure 7. The variance component parameter* $\tau = 0.1$ *and 0.25. Numbers were presented in Supplementary Table 4. B. based on 8.3x10*$^{8}$ *association tests. or each censoring rate, 100 data sets were simulated. Each data set contains 10,000 randomly selected individuals with white British ancestry. The variance component parameter* $\tau = 0.25$*. Numbers were presented in Supplementary Table 5.*

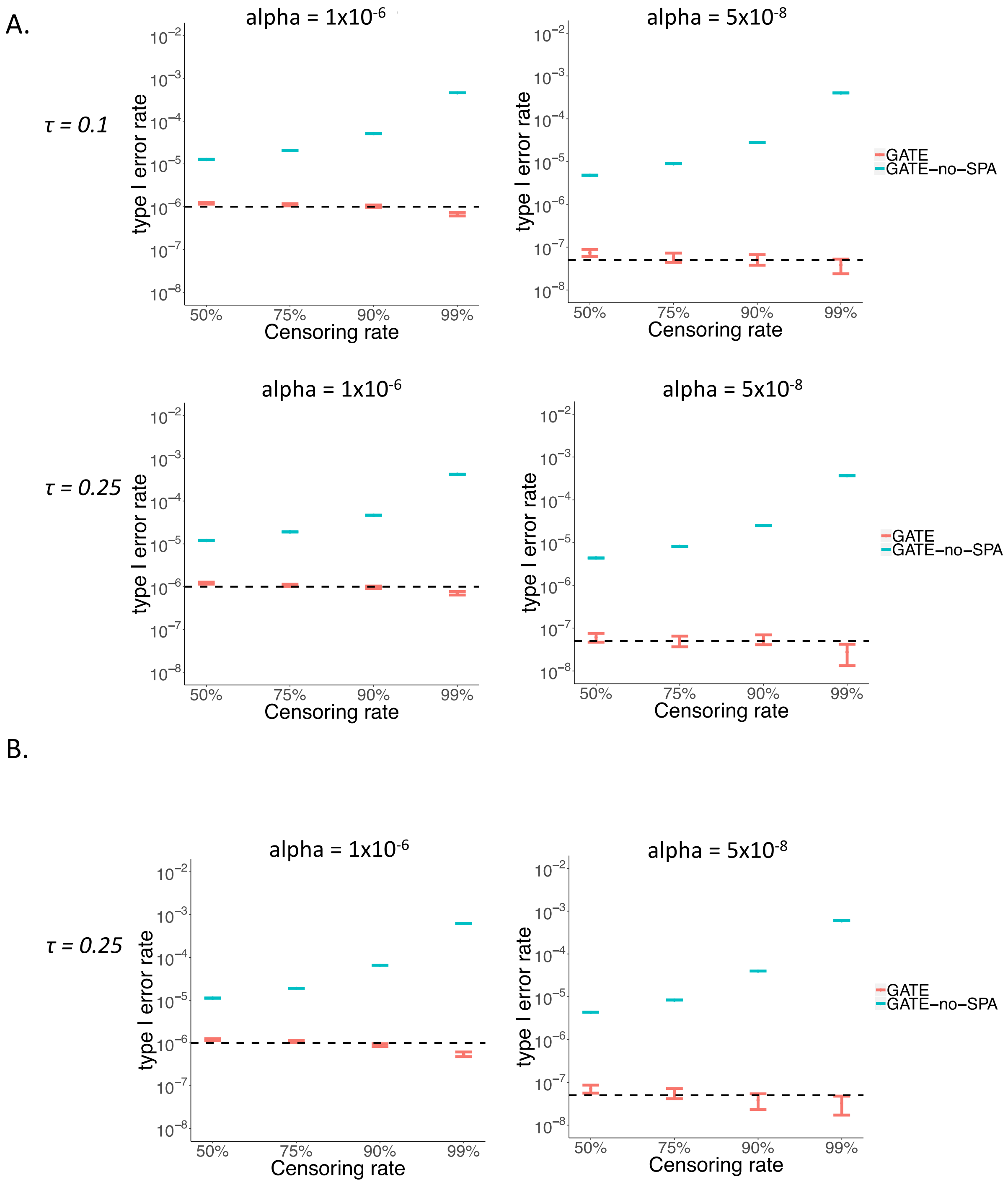

*Supplementary Figure 9: Quantile-quantile plots stratified by MAF for randomly selected 10 million association tests from the simulation study for evaluating type I error rates using10,000 randomly selected individuals with white British ancestry (Supplementary Table 5).*

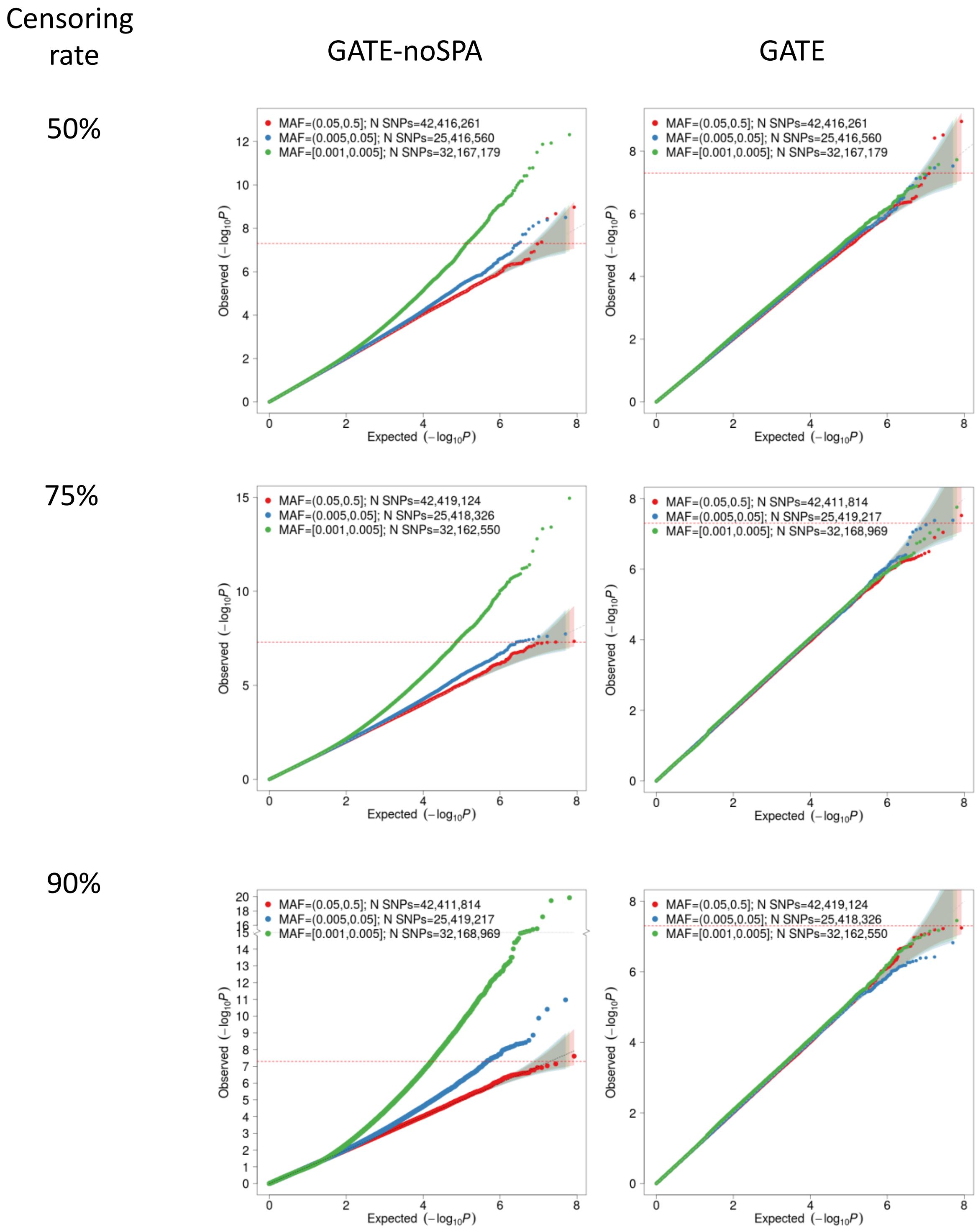

*Supplementary Figure 10: Empirical power of GATE and COXMEG at the significance level α = 5×10^-8, when the variance component parameter τ = 0.25 and the censoring rate is 50%.*

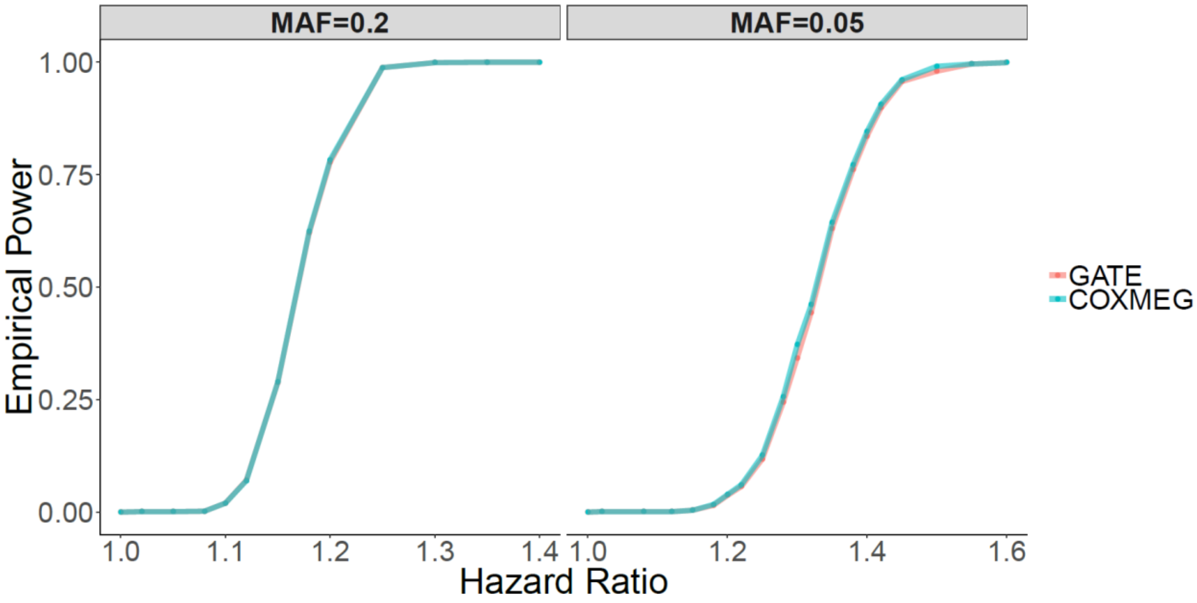

*Supplementary Figure 11: Comparison of GATE association p-values between using different time-units for defining the event times of the four time-to-event phenotypes based on the UK Biobank data. Results were compared among the event and censoring times specified in nearest 1 month, 3 months, 6 months and 12 months time-units. Four phenotypes (ischemic heart disease, female breast cancer, glaucoma, and Alzheimer's disease) were analysed for association across 46 million imputed genetic variants with INFO ≥ 0.3 and MAC ≥ 20.*

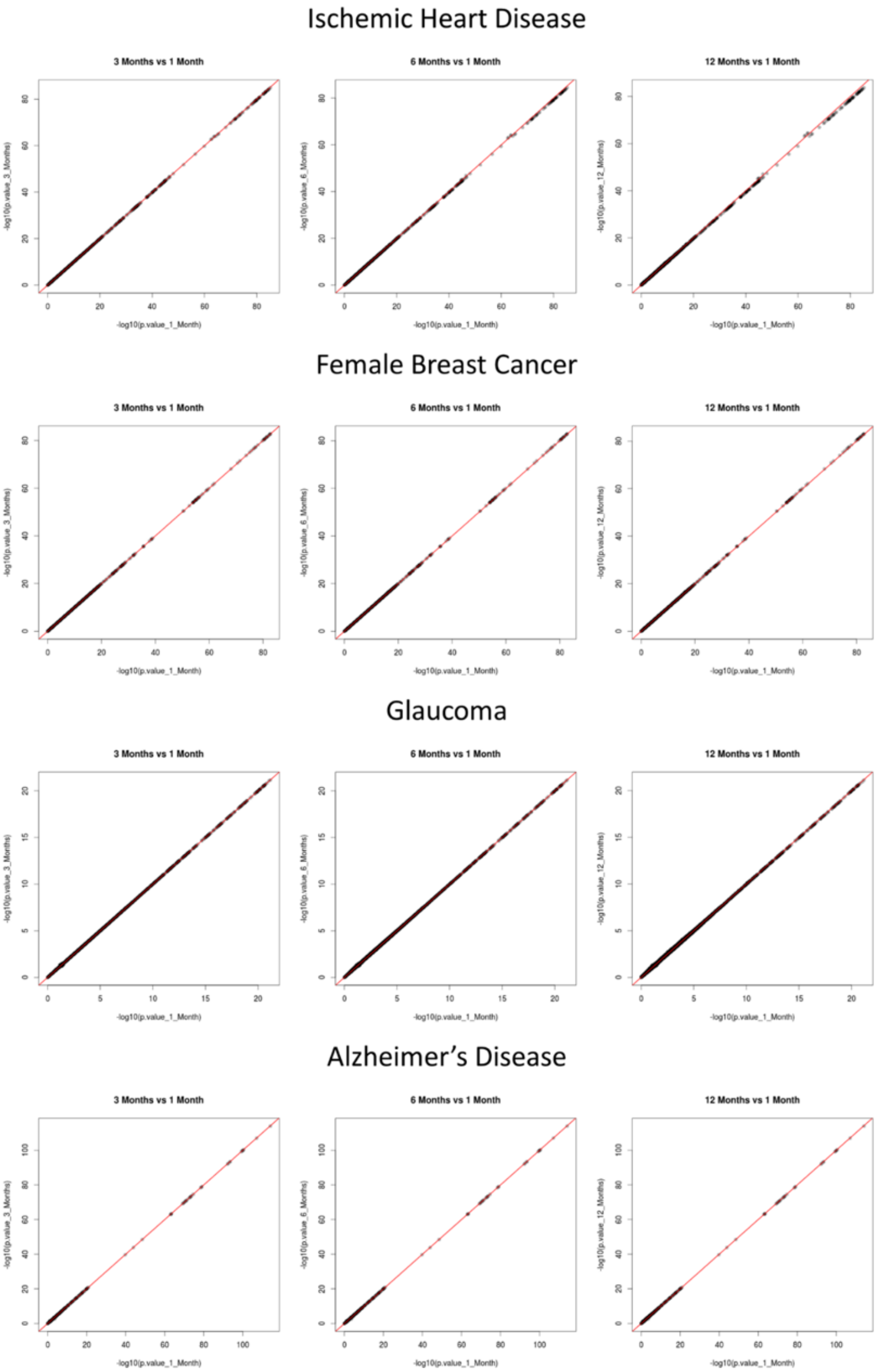

*Supplementary Figure 12: Comparison of association p-values between using different number of markers for constructing the GRM in step 1 of GATE for analysing the four time-to-event phenotypes based on the UK Biobank data. Results were compared between 93511 high-quality genotyped markers used by the UK Biobank research group for estimating kinship, and 245745 pruned markers with MAF>1%. Four phenotypes (ischemic heart disease, female breast cancer, glaucoma, and Alzheimer's disease) were analysed for association across 46 million imputed genetic variants with INFO ≥ 0.3 and MAC ≥2 0.*

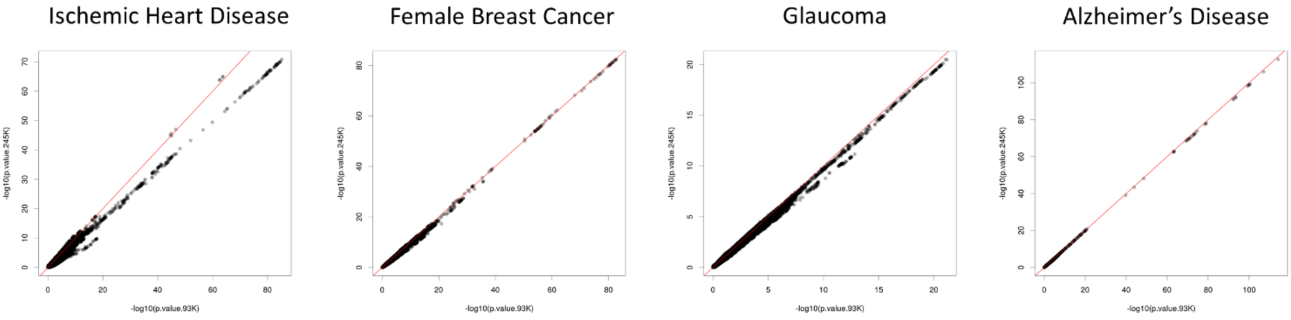


*Supplementary Figure 13: Manhattan plots for GWAS of four binary phenotypes based the UK Biobank subjects with White British ancestry using GATE with A) 93,511 markers used in the GRM, and B) 245,745 markers used in the GRM. Four phenotypes (ischemic heart disease, female breast cancer, glaucoma, and Alzheimer's disease) were analysed for association across 46 million imputed genetic variants with INFO ≥ 0.3 and MAC ≥ 20.*

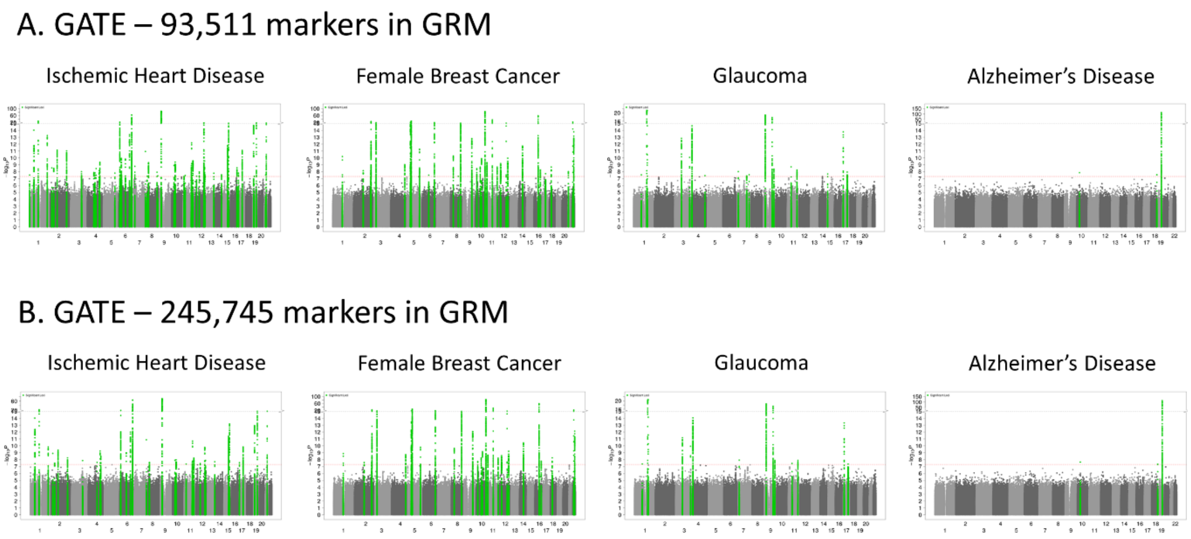


*Supplementary Figure 14: Estimated variance ratios using different number of randomly selected markers for four phenotypes based on the UK Biobank subjects with White British ancestry using GATE.*

*For each choice of number of markers, we randomly selected the markers 50 times. The dotted lines represent the variance ratios with 500 randomly selected markers.*

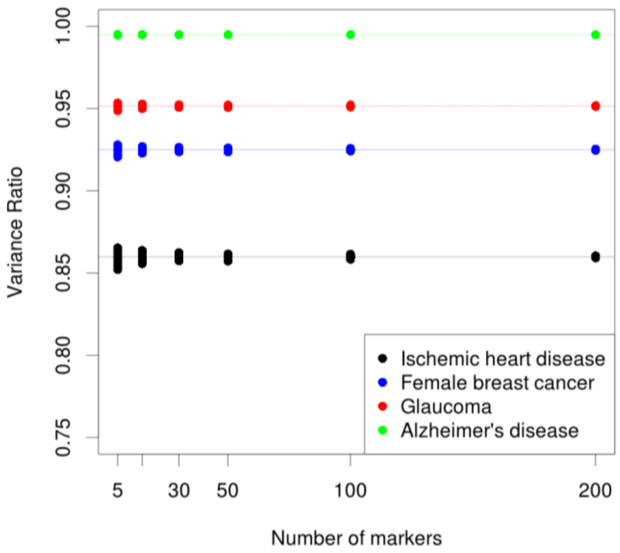
