## Supplementary Notes for "An efficient and accurate frailty model approach for genome-wide survival association analysis controlling for population structure and relatedness in large-scale biobanks"

### 1 Derivation of the GATE Method

#### 1.1 Derivation of the likelihood

Consider a set of $N$ subjects where $E_i$ and $C_i$ denote the failure and censoring times respectively, for subject $i = 1, \ldots, N$. Let $T_i = \min(E_i, C_i)$, and $\Delta_i = I(E_i \leq C_i)$, where $I(.)$ denotes the indicator function. For each subject, we have the observed event status-time pair $(T_i = t_i, \Delta_i = \delta_i)$. We call $t_i$ a failure time for subject $i$ if $\delta_i = 1$, and a censoring time if $\delta_i = 0$. Without loss of generality, we assume $t_i$s are in increasing order, i.e, $t_1 \leq t_2 \leq \ldots \leq t_N$.

Let $X_i$ denote the $q \times 1$ vector of covariates (excluding the intercept) to adjust for, and $G_i$ denote the genotype $(0, 1, \text{ or } 2)$ of the variant we are testing. Throughout this paper, quantities without a subscript is used to denote the vectors of corresponding quantities with subscripts (eg. $G = (G_1, \ldots, G_N)$), when it is clear from the context. Denote $X$ to be the $n \times q$ covariate matrix with $i$-th row as $X_i^\top$. Then, in a frailty model, we denote the conditional hazard function for subject $i$ at time $t$ given the random effects $b_i$s,

$$\lambda_i(t|b) = \lambda_0(t) \exp(\eta_i); \qquad \eta_i = X_i^\top \beta + G_i \gamma + b_i,$$

and the survival function $S_i(t|b) = \exp(-\Lambda_0(t) \exp(\eta_i))$, where $\Lambda_0(t) = \int_0^t \lambda_0(u) du$ is the cumulative baseline hazard (CBH) function. Under multivariate Gaussian frailty, the random effects $b \sim N(0, \tau V)$, where $\tau$ is the variance component, and $V$ is the $N \times N$ genetic relationship matrix (GRM). We further assume that conditional on $b$, the censoring is independent and non-informative of $b$. Then the likelihood of the observed data is,

$$\begin{aligned} L(\lambda_0, \beta, \gamma, \tau) &= \int \prod_{i=1}^{N} \left[\lambda_i(t_i|b)^{\delta_i} S_i(t_i|b)\right] \frac{1}{|\tau V|^{1/2}} \exp\left(-\frac{1}{2} b^\top (\tau V)^{-1} b\right) db \\ &= \frac{1}{|\tau V|^{1/2}} \prod_{i=1}^{N} \lambda_0(t_i)^{\delta_i} \int \exp\left[\sum_{i=1}^{N} (\delta_i \eta_i - \Lambda_0(t_i) \exp(\eta_i)) - \frac{1}{2} b^\top (\tau V)^{-1} b\right] db. \end{aligned}$$

Let $f(b)=\sum_{i=1}^N\left(\delta_i\eta_i-\Lambda_0(t_i)\exp(\eta_i)\right)-\frac{1}{2}b^\top(\tau V)^{-1}b$. Then we can approximate the integral using the Laplace approximation,

$$\int \exp f(b)db \approx (2\pi)^{N/2}\left|-f''(\tilde{b})\right|^{-1/2}\exp(f(\tilde{b})),$$

where $\tilde{b}=\arg_b\max f(b)$ is the solution to $f'(b)=0$. Therefore, the log-likelihood can be approximated by,

$$\begin{aligned}
\ell((\lambda_0,\beta,\gamma,\tau) \approx & -\frac{1}{2}\log|\tau V|+\sum_{i=1}^N\delta_i\log\lambda_0(t_i)-\frac{1}{2}\log\left|-f''(\tilde{b})\right|+f(\tilde{b}) \\
= & -\frac{1}{2}\log|\tau V|-\frac{1}{2}\log\left|\sum_{i=1}^N\Lambda_0(t_i)\exp(\tilde{\eta}_i)+(\tau V)^{-1}\right| \\
& +\sum_{i=1}^N\left[\delta_i\left(\log\lambda_0(t_i)+\tilde{\eta}_i\right)-\Lambda_0(t_i)\exp(\tilde{\eta}_i)\right]-\frac{1}{2}\tilde{b}^\top(\tau V)^{-1}\tilde{b} \qquad (1) \\
= & -\frac{1}{2}\log|\tau V|-\frac{1}{2}\log\left|\tilde{W}+(\tau V)^{-1}\right| \\
& +\sum_{i=1}^N\left[\delta_i\left(\log\lambda_0(t_i)+\tilde{\eta}_i\right)-\tilde{\mu}_i\right]-\frac{1}{2}\tilde{b}^\top(\tau V)^{-1}\tilde{b},
\end{aligned}$$

where $\tilde{\eta}_i=X_i^\top\beta+G_i\gamma+\tilde{b}_i$, $\tilde{\mu}_i=\Lambda_0(t_i)\exp(\tilde{\eta}_i)$, and $\tilde{W}=diag(\tilde{\mu}_i)$. The log-likelihood (1) with respect to $\beta$ and $\tau$ is similar to the Poisson GLMM log-likelihood.[1,2] Following Breslow and Clayton,[2] assuming the weight matrix $W$ changes slowly as a function of the mean, we can maximize the following penalized log-likelihood[3] given $\tau$,

$$\ell_p\left(\lambda_0,\beta,\gamma,b\right)=\sum_{i=1}^N\left[\delta_i\left(\log\lambda_0(t_i)+\eta_i\right)-\mu_i\right]-\frac{1}{2}b^\top(\tau V)^{-1}b, \qquad (2)$$

to obtain the maximum likelihood estimators (MLEs) $(\hat{\lambda}_0,\hat{\beta},\hat{\gamma},\hat{b})=(\hat{\lambda}_0^\tau,\hat{\beta}^\tau,\hat{\gamma}^\tau,\hat{b}^\tau)$, where $\hat{b}^\tau=\tilde{b}(\hat{\beta}^\tau)$, and $\mu_i=\Lambda_0(t_i)\exp(\eta_i)$.

#### 1.2 Estimation of the baseline hazard, fixed effects and random effects given the variance component

Lets denote the unique failure times to be $t_1^*<t_2^*<\ldots<t_K^*$ in increasing order. Following Breslow's suggestion,[4] we assume the baseline hazard function to remain constant between successive event times, and according to Kalbfleisch and Prentice's[5] convention, we consider the censored observations to be censored at the previous failure time. In section 6 we show that the maximum likelihood estimation under these assumptions is equivalent to maximizing a partial likelihood.[6] Formally, lets denote $\lambda_0(t)=\alpha_i$, for $t_i^*\le t<t_{i+1}^*$. Then the cumulative hazard function can be written as a step function,

$$\Lambda_0(t)=\sum_{j=1}^i\alpha_j\left(t_j^*-t_{j-1}^*\right),\quad t_i^*\le t<t_{i+1}^*,$$

with the convention $t_0^* = t_0 = 0$. Then, using algebraic manipulations with the order of the summations, we can write,

$$\sum_{i=1}^{N} \mu_i = \sum_{i=1}^{K} \left[ \alpha_i (t_i^* - t_{i-1}^*) \sum_{j \in R(t_i^*)} \exp(\eta_j) \right],$$

where $R(t_i) = \{j : t_j \geq t_i\}$ denotes the set of subjects at risk at time $t_i$. This definition of the at-risk set corresponds to Breslow's approximation when there are tied failure times.[7] We can express $\ell_p$ as a function of $(\alpha, \beta, \gamma, b)$, and obtain the score functions by taking derivatives of $\ell_p$ with respect to $\beta, \gamma, b$, and $\alpha_i$-s,

$$\begin{aligned} \frac{\partial \ell_p}{\partial \beta} &= \sum_{i=1}^{N} (\delta_i - \mu_i) X_i \\ \frac{\partial \ell_p}{\partial \gamma} &= \sum_{i=1}^{N} (\delta_i - \mu_i) G_i \\ \frac{\partial \ell_p}{\partial b} &= \sum_{i=1}^{N} (\delta_i - \mu_i) Z_i - (\tau V)^{-1} b \\ \frac{\partial \ell_p}{\partial \alpha_i} &= \frac{d_i}{\alpha_i} - (t_i^* - t_{i-1}^*) \sum_{j \in R(t_i^*)} \exp(\eta_j), \end{aligned} \tag{3}$$

where $Z_i$ is an $N \times 1$ vector with $i$-th element 1 and rest 0s, and $d_i = \sum_{j: t_j = t_i^*} \delta_j$ is the number of failures at time $t_i^*$. Setting $\partial \ell_p / \partial \alpha_i = 0$, we obtain the MLEs,

$$\hat{\alpha}_i = d_i \left[ (t_i^* - t_{i-1}^*) \sum_{j \in R(t_i^*)} \exp(\eta_j) \right]^{-1}.$$

This leads to the famous Breslow's estimator for the CBH,

$$\hat{\Lambda}_0(t) = \sum_{i=1}^{N} \frac{\delta_i I(t_i \leq t)}{\sum_{j \in R(t_i)} \exp(\eta_j)}.$$

For the score test of $H_0 : \gamma = 0$ vs $H_1 : \gamma \neq 0$, we only estimate the MLEs $(\hat{\alpha}, \hat{\beta}, \hat{b})$ under $H_0$, setting $\gamma = 0$. Using similar techniques to estimate fixed effects parameters in Poisson GLMM, we denote the working outcome vector $Y = \eta + W^{-1}(\delta - \mu)$, where $W = diag(\mu_i)$, $\eta = (\eta_1, \ldots, \eta_N)$, $\delta = (\delta_1, \ldots, \delta_N)$, and $\mu = (\mu_1, \ldots, \mu_N)$. Then, $\delta - \mu = W(Y - \eta) = W(Y - X\beta - Zb)$, and the score equations can be written as,

$$\begin{bmatrix} X^\top W X & X^\top W \\ WX & W + (\tau V)^{-1} \end{bmatrix} \begin{bmatrix} \beta \\ b \end{bmatrix} = \begin{bmatrix} X^\top W Y \\ WY \end{bmatrix}$$

Let $\Sigma = W^{-1} + \tau V$, then

$$\begin{aligned} \hat{\beta} &= \left( X^\top \Sigma^{-1} X \right)^{-1} X^\top \Sigma^{-1} Y \\ \hat{b} &= \tau V \Sigma^{-1} \left( Y - X\hat{\beta} \right). \end{aligned}$$

##### 1.2.1 Estimation of the variance component

Given $\hat{\alpha} = \hat{\alpha}(\tau), \hat{\beta} = \hat{\beta}(\tau)$ estimated, from (1) the log-likelihood of the variance component can be derived as,

$$\ell(\hat{\alpha}(\tau), \hat{\beta}(\tau), \gamma = 0, \tau) = -\frac{1}{2}log|\Sigma| - \frac{1}{2}Y^\top PY,$$

where $P = \Sigma^{-1} - \Sigma^\top X \left(X^\top \Sigma^{-1} X\right)^{-1} X^\top \Sigma^{-1}$. We maximize the corresponding restricted maximum-likelihood (REML),

$$\ell_R(\hat{\alpha}(\tau), \hat{\beta}(\tau), \gamma = 0, \tau) = -\frac{1}{2}log|\Sigma| - \frac{1}{2}log|X^\top \Sigma^{-1} X| - \frac{1}{2}Y^\top PY.$$

The score function with respect to $\tau$ are given by,

$$U_\tau = \frac{\partial \ell_R(\hat{\alpha}(\tau), \hat{\beta}(\tau), \gamma = 0, \tau)}{\partial \tau} = \frac{1}{2}\left[Y^\top PVPY - tr(PV)\right].$$

The corresponding observed information function, and the expected information function are given by

$$J_\tau = -\frac{\partial^2 \ell_R(\hat{\alpha}(\tau), \hat{\beta}(\tau), \gamma = 0, \tau)}{\partial \tau^2} = -\frac{1}{2}tr(PVPV) + Y^\top PVPVPY,$$
$$E\left(J_\tau\right) = E\left[-\frac{\partial^2 \ell_R(\hat{\alpha}(\tau), \hat{\beta}(\tau), \gamma = 0, \tau)}{\partial \tau^2}\right] = \frac{1}{2}tr(PVPV),$$

respectively. Evaluating both observed and expected information functions involves computationally expensive trace computations. To avoid the trace computations, the average information is used in the AI-REML[1,8,9] algorithm. The average information is expressed as the average of $J_\tau$ and $E(J_\tau)$,

$$AI_\tau = \frac{1}{2}Y^\top PVPVPY.$$

##### 1.2.2 Algorithm to fit the null model

The null model fitting algorithm can be summarized as,

1. Fit a Poisson linear model wth $\tau = 0$ to get initial estimates $\hat{\beta}^{(0)}$ and working outcome vector $Y^{(0)}$.
2. Calculate Breslow's estimator $\hat{\Lambda}_0^{(0)}(t)$.
3. At the $i$-th step, update $\hat{\tau}$ using $\hat{\tau}^{(i)} = \hat{\tau}^{(i-1)} + \left\{AI_\tau\Big|_{\tau=\hat{\tau}^{(i-1)}}\right\}^{-1} \left\{U_\tau\Big|_{\tau=\hat{\tau}^{(i-1)}}\right\}$.
4. Update $\hat{\beta}, \hat{b}, \hat{\alpha}$ using $Y$ and $\hat{\tau}^{(i)}$.
5. Update $Y$ and $\hat{\Lambda}_0(t)$ using $\hat{\beta}^{(i)}, \hat{b}^{(i)}, \hat{\alpha}^{(i)}, \hat{\tau}^{(i)}$.
6. Repeat steps 2–5, until $\max\left\{\frac{\left|\hat{\beta}^{(i)} - \hat{\beta}^{(i-1)}\right|}{\left|\hat{\beta}^{(i)}\right| + \left|\hat{\beta}^{(i-1)}\right|}, \frac{\left|\hat{\tau}^{(i)} - \hat{\tau}^{(i-1)}\right|}{\left|\hat{\tau}^{(i)}\right| + \left|\hat{\tau}^{(i-1)}\right|}\right\} \leq tolerance$.

#### 1.3 Score test

The score test statistic under the null hypothesis is given by,

$$T = \frac{\partial l}{\partial \gamma}\Big|_{\left(\hat{\beta},\hat{b},\gamma=0,\hat{\alpha},\hat{\tau}\right)} = G^\top(\delta - \hat{\mu}) = \tilde{G}^\top(\delta - \hat{\mu}),$$

where $\tilde{G} = G - \tilde{X}\left(\tilde{X}^\top \hat{W} \tilde{X}\right)^{-1} \tilde{X}^\top \hat{W} G$ is the covariate-and-intercept adjusted genotype vector, and $\tilde{X} = \begin{bmatrix} 1 & X \end{bmatrix}$ is the intercept-augmented covariate matrix. The information matrix corresponding to the score equations in (3) is given by,

$$I(\beta,\gamma,b,\alpha) = \begin{bmatrix} X^\top W X & X^\top W G & X^\top W & B^{(X)\top} \\ G^\top W X & G^\top W G & G^\top W & B^{(G)\top} \\ W X & Z^\top W G & W + (\tau V)^{-1} & B^{(Z)\top} \\ B^{(X)} & B^{(G)} & B^{(Z)} & B^{(\lambda)} \end{bmatrix},$$

where,

$$\begin{aligned} B^{(\lambda)} &= diag\left(-\frac{\partial^2 \ell_p}{\partial \alpha_i}\right) = diag\left(d_i/\alpha_i^2\right) \\ B_{\bullet i}^{(X)\top} &= -\frac{\partial^2 \ell_p}{\partial \alpha_i \partial \beta^\top} = (t_i^* - t_{i-1}^*) \sum_{j\in R(t_i^*)} \exp(\eta_j) X_j \\ B_{\bullet i}^{(G)\top} &= -\frac{\partial^2 \ell_p}{\partial \alpha_i \partial \gamma^\top} = (t_i^* - t_{i-1}^*) \sum_{j\in R(t_i^*)} \exp(\eta_j) G_j \\ B_{\bullet i}^{(Z)\top} &= -\frac{\partial^2 \ell_p}{\partial \alpha_i \partial b^\top} = (t_i^* - t_{i-1}^*) \sum_{j\in R(t_i^*)} \exp(\eta_j) Z_j. \end{aligned}$$

$A_{ij}$ and $A_{\bullet i}$ denote the $(i,j)$-th element and $i$-th column of a matrix $A$, respectively. Then, the variance of the score statistic under $H_0$ is given by,

$$Var_{H_0}(T) = \left(I(\hat{\beta},\hat{b},\gamma=0,\hat{\alpha})^{-1}\right)_{22} = G^\top \hat{Q} G = \tilde{G}^\top \hat{Q} \tilde{G},$$

where $\hat{Q} = \hat{S}^{-1} - \hat{S}^{-1} X \left(X^\top S^{-1} X\right)^{-1} X^\top \hat{S}^{-1}$, $\hat{S} = (\hat{W} - \hat{U})^{-1} + \hat{\tau} V$. The matrix $\hat{U}$ is defined as,

$$\hat{U} = \sum_{i=1}^{K} \frac{d_i}{(v_i^\top 1)^2} v_i v_i^\top = \Gamma R^\top D R \Gamma,$$

$$D = diag\left\{\frac{d_i}{\left(\sum_{j\in R(t_i^*)} \exp(\hat{\eta}_j)\right)^2}\right\}_{i=1}^{K}, \qquad \Gamma = diag\left\{\exp(\hat{\eta}_i)\right\}_{i=1}^{N},$$

$$R_{ij} = \begin{cases} 0, & \text{if } j \notin R(t_i^*) \\ 1, & \text{if } j \in R(t_i^*) \end{cases}, \quad \text{for } i = 1,\dots,K, j = 1,\dots,N,$$

where $v_i$ is an $n$-dimensional vector with elements $v_{ij} = 0$ if $j \notin R(t_i^*)$, and $v_{ij} = \exp(\hat{\eta}_j)$ if $j \in R(t_i^*)$.

#### 2 Approaches to reduce computation and memory cost

To obtain $Var_{H_0}(T) = \tilde{G}^\top \hat{Q} \tilde{G}$, we need to compute quantities of the form $\hat{S}^{-1}a$, where $a$ is a vector. The standard computation technique of inverting $\hat{S}$ (computation cost $O(N^3)$) and multiplying $\hat{S}^{-1}$ with $a$ can be extremely time consuming when $N$ is large. On top of that, it will require the storage of $\hat{S}^{-1}$ (memory cost $O(N^2)$) which can have extremely high memory requirement. In order to reduce the computation and memory cost, we implemented several strategies similar to BOLT-LMM[10] and SAIGE.[9] Firstly, instead of storing the $N \times N$ GRM which can cost $4N(N+1)$ bytes if stored using double precision floating point numbers, we only store the raw genotypes as binary vectors which only costs $NM/4$ bytes, where $M$ is the number of markers used to calculate the GRM. We calculate the elements of the GRM from the raw genotype vectors only when they are needed. Secondly, to compute quantities of the form $\hat{S}^{-1}a$, we implemented the pre-conditioned conjugate gradient[11] (PCG) method, which computes $\hat{S}^{-1}a = b$ by solving the linear system of equations $\hat{S}b = a$. Thirdly, since $U = \Gamma R^\top D R \Gamma$ has a low rank decomposition with $rank(U) = K$, the number of unique failure times, to calculate $(W-U)^{-1}G$ in the PCG steps, we leverage the Woodbury identity decomposition

$$(W-U)^{-1}G = W^{-1}G - W^{-1}\Gamma R^T(-D^{-1} + R\Gamma W^{-1}\Gamma R^T)^{-1} R\Gamma W^{-1}G.$$

The matrix $(-D^{-1} + R\Gamma W^{-1}\Gamma R^T)$ is a $K \times K$ matrix which can be inverted easily when $K$ is small, or when $K$ is also larger, PCG can be used to obtain $(-D^{-1} + R\Gamma W^{-1}\Gamma R^T)^{-1} R\Gamma W^{-1}G$.

Similar to SAIGE, we further implemented the Hutchinson's randomized trace estimation[12,13] for calculating $tr(PV)$. We also implemented multithreaded parallel computation for the matrix-vector multiplications in the PCG steps using Intel Threading Building Block (TBB) from the RcppParallel[14] package.

#### 3 Variance ratio approximation

Computation of the variance of the score statistic $Var_{H_0}(T) = \tilde{G}^\top \hat{Q} \tilde{G}$ requires calculating $\hat{Q}\tilde{G}$ repeatedly for all markers, which is computationally expensive. To avoid calculating $\hat{Q}\tilde{G}$ for all the markers, we estimate the variance ratio $\hat{r} = \tilde{G}^\top \hat{Q} \tilde{G} / \tilde{G}^\top \hat{W} \tilde{G}$ using a small set of markers, and then approximate the variance of the score statistic for all markers by $\hat{r}\tilde{G}^\top \hat{W} \tilde{G}$ in step 2. This saves substantial computation time since $\hat{W}$ is a diagonal matrix. Such variance ratio approximation approaches were previously used in various linear[10,15,16] and logistic mixed effects models[9] to speed up computation. Here, we provide a theoretical justification for why such an approximation works.

Let $P_{\tilde{X}} = \tilde{X}(\tilde{X}^\top \hat{W} \tilde{X})^{-1} \tilde{X}^\top \hat{W}$ be the weighted projection matrix for the intercept-augmented covariate matrix $\tilde{X}$. Let $E(G_i) = \mu_g$ and the covariance matrix of $G$ is given by $\sigma_g^2 \Psi$, where $\Psi$ is the correlation matrix of $G$. When the elements of $G$ follows the $Bin(2, p_g)$ distribution, then $\mu_g = 2p_g$, and $\sigma_g^2 = \sqrt{2p_g(1-p_g)}$. The matrix $\Psi$ represents the kinship matrix, however exact characterization of $\Psi$ is not needed for this proof. Then, $E(\tilde{G}) = \mu_g(I - P_{\tilde{X}})1 = 0$, and $Cov(G) = \sigma_g^2 (I - P_{\tilde{X}}) \Psi (I - P_{\tilde{X}})^\top$, where 1 is the $N \times N$ vector of all element equal to unity. We scale both the numerator and denominator of the variance ratio by $N^{-1}$ so that they don't blow to infinity when looked at individually. Then, for the numerator,

$$E(N^{-1}\tilde{G}^\top \hat{Q}\tilde{G}) = \frac{\sigma_g^2}{N} tr\left[\hat{Q}(I - P_{\tilde{X}})\Psi(I - P_{\tilde{X}})^\top\right] = \frac{\sigma_g^2}{N} tr(\hat{Q}\Psi),$$

since $(I - P_{\tilde{X}})^\top \hat{Q}(I - P_{\tilde{X}}) = \hat{Q}$. Similarly, for the denominator,

$$E(N^{-1}\tilde{G}^\top \hat{W}\tilde{G}) = \frac{\sigma_g^2}{N} tr\left[\hat{W}(I - P_{\tilde{X}})\Psi(I - P_{\tilde{X}})^\top\right] = \frac{\sigma_g^2}{N} tr\left[\hat{W}(I - P_{\tilde{X}})\Psi\right],$$

since $(I - P_{\tilde{X}})^\top \hat{W}(I - P_{\tilde{X}}) = \hat{W}(I - P_{\tilde{X}})$. Therefore, as the eigenvalues of $\hat{Q}, \hat{W}, \Psi$ are bounded, and the distribution of $G$ has bounded support, the variances of the numerator and the denominator terms are both $O(N^{-1})$, and the variance ratio converges to,

$$\hat{r} = \frac{\tilde{G}^\top \hat{Q}\tilde{G}}{\tilde{G}^\top \hat{W}\tilde{G}} \xrightarrow{p} \frac{lim_{N\to\infty}\left\{N^{-1}tr(\hat{Q}\Psi)\right\}}{lim_{N\to\infty}\left\{N^{-1}tr\left[\hat{W}(I - P_{\tilde{X}})\Psi\right]\right\}}.$$

The ratio on the right-hand side is constant across all markers as the individual limits in the numerator and denominator exist and are bounded away from zero.

Our software first estimates the variance ratio using 30 randomly selected markers, and then adds more markers with increments of ten until the coefficient of variation (CV) is smaller than 0.001. We performed a sensitivity analysis on the number of markers used to calculate the variance ratio using the UK Biobank data on white British subjects. For the analysis of the four exemplary phenotypes described in this paper, we used different number of markers ($N = 5, 15, 30, 50, 100, 200$) to estimate the variance ratios. For each choice of number of markers, we selected the markers randomly 50 times and the variance ratio estimates are presented in Supplementary Figure 14. The results show that the variance ratio estimates remain overall very stable when $\geq 30$ markers are used, and the variation decreases with increasing number of markers.

### 4 Using saddlepoint approximation[17] (SPA) for the null distribution of the score statistic

In traditional score tests, the distribution of the score statistic under $H_0$ is approximated by a Normal distribution, which uses the first two moments, mean and variance. This approach can perform poorly in the tail regions, especially if the underlying distribution is highly skewed when the studying event is very rare or the testing genetic variant has a very low minor allele count (MAC). Here, similar to what has been applied in the logistic mixed models[9,18–20] previously, we use the SPA to approximate the null distribution of the score statistic to obtain accurate p-values. Based on the likelihood (1), we derive the SPA of $T_{adj} = \tilde{G}^\top(\delta - \hat{\mu})/\sqrt{\hat{r}\tilde{G}^\top \hat{W}\tilde{G}}$ as a weighted sum of independent Poisson random variables given $b$. The approximated cumulant generating function (CGF) of $T_{adj}$ and it's derivatives are given by,

$$K(\xi;\hat{\mu}) = \sum_{i=1}^{N} \hat{\mu}_i(e^{\tilde{G}_i c\xi} - \tilde{G}_i c\xi - 1),$$
$$K'(\xi;\hat{\mu}) = \sum_{i=1}^{N} \hat{\mu}_i \tilde{G}_i c(e^{\tilde{G}_i c\xi} - 1),$$
$$K''(\xi;\hat{\mu}) = \sum_{i=1}^{N} \hat{\mu}_i \tilde{G}_i^2 c^2 e^{\tilde{G}_i c\xi},$$

where $c = (\tilde{G}^\top W \tilde{G})^{-1/2}$. To calculate the probability that $T_{adj} < s$, where $s$ is the observed test statistic, we use the following formula

$$Pr(T < s) = \Phi\left\{w + \frac{1}{w} log\left(\frac{v}{w}\right)\right\},$$

where $w = sign(\hat{\xi})[2\{\hat{\xi}s - K(\hat{\xi})\}]^{\frac{1}{2}}$, $v = \hat{\xi}[K''(\hat{\xi})]^{\frac{1}{2}}$, $\hat{\xi}$ is the solution of $K'(\hat{\xi}) = s$, and $\Phi$ is the standard normal distribution function.

### 5 Effect size estimation

Since our method only fits the model under the null hypothesis of no association, it cannot provide the effect size estimates as part of the model fitting process. Instead, to rapidly estimate the effect sizes, we follow a similar approach used in EMMAX,[21] GRAMMAR-Gamma,[15] and SAIGE[9] using the parameter estimates from the null model. Our genetic effect size estimate is given by $\hat{\gamma} = \left(\tilde{G}^\top \hat{Q} \tilde{G}\right)^{-1} \tilde{G}^\top \hat{Q}(\delta - \hat{\mu})$. Notice that this estimate can also be expressed as $\hat{\gamma} = T/Var_{H_0}(T)$ where $T$ is the score test statistic, and derived assuming the standardized Wald test statistic to be equal to the standardized score test statistic. In section 3, we showed that $Var_{H_0}(T) = \tilde{G}^\top \hat{Q} \tilde{G} \approx \hat{r}\tilde{G}^\top \hat{W} \tilde{G}$. Therefore, $\hat{\gamma} = T/\hat{r}\tilde{G}^\top \hat{W} \tilde{G}$, which can be estimated using the already estimated quantities $T$,$\hat{r}$, and $\tilde{G}^\top \hat{W} \tilde{G}$. The standard errors can be estimated by inverting the p-values. The standard error of $\hat{\gamma}$, $SE(\hat{\gamma}) = |\hat{\gamma}/z|$, where $z$ is the Z-score corresponding to the two-sided association p-value.

### 6 Equivalence of penalized full and partial likelihood-based estimates

For unrelated samples, the equivalence between maximizing the partial likelihood, and maximizing the full likelihood with Breslow's estimator plugged in as the estimate of the CBH function has been shown by Breslow[4] in the proportional hazard model. Here, we show that given the variance component $\tau$, maximizing $\ell_p$ (see (2)) assuming that the baseline hazard to be constant between any two consecutive failure times, i.e., estimating the CBH function using Breslow's estimator, is equivalent to maximizing the penalized partial log-likelihood as described in Ripati et al.,[6]

$$\ell_{pp}(\beta,\gamma,b) = \sum_{i=1}^{N} \delta_i \left( \eta_i - \log \sum_{j \in R(t_i)} \exp(\eta_j) \right) - \frac{1}{2} b^\top (\tau V)^{-1} b. \tag{4}$$

Plugging in $\hat{\Lambda}_0(t)$ into the score equations corresponding to $\beta, \gamma$ in (3), and using algebraic manipulations with the ordering of the summations, we get,

$$\frac{\partial \ell_p}{\partial \beta} = \sum_{i=1}^{n} \delta_i \left( X_i - \frac{\sum_{j \in R(t_i)} \exp(\eta_j) X_j}{\sum_{j \in R(t_i)} \exp(\eta_j)} \right),$$

$$\frac{\partial \ell_p}{\partial \gamma} = \sum_{i=1}^{n} \delta_i \left( G_i - \frac{\sum_{j \in R(t_i)} \exp(\eta_j) G_j}{\sum_{j \in R(t_i)} \exp(\eta_j)} \right).$$

We observe that the expressions of the score functions are the same as the score functions from the partial likelihood $\partial \ell_{pp}/\partial \beta$,$\partial \ell_{pp}/\partial \gamma$. Equivalence of the information matrices can also be shown similarly.

### 7 Sensitivity analysis on the number of markers used to construct the GRM

We performed a sensitivity analysis on the number of markers used to construct the GRM in step 1 of GATE. We compared association results between using 93511 high-quality genotyped markers that were used by the UK Biobank research group to calculate kinship,[22] and 245745 pruned (500kb window, sliding step-size 50 markers, $r^2 < 0.2$) genotyped markers with MAF $\geq 0.01$. We compared the association results between these two marker-sets for the analysis of four example phenotypes in the UK Biobank data on 46 million imputed variants. Manhattan plots (Supplementary Figure 13) were similar between the two marker-sets, and the scatter plots of the p-values (Supplementary Figure 12) show highly correlated association p-values. P-values using 245745 markers were generally slightly smaller than the p-values using 93511 markers, especially for ischemic heart disease. Similar observation was also made for the analysis of the binary disease status for this phenotype.[9]

[5] J. D Kalbfleisch. *The statistical analysis of failure time data.* Wiley series in probability and mathematical statistics. Wiley, New York, 1980.

[6] Samuli Ripatti and Juni Palmgren. Estimation of multivariate frailty models using penalized partial likelihood. *Biometrics*, 56(4):1016–1022, 2000.

[7] Terry M. Therneau. *Modeling Survival Data: Extending the Cox Model.* Statistics for Biology and Health. Springer New York : Springer, New York, NY, 2000.

[14] JJ Allaire, Romain Francois, Kevin Ushey, Gregory Vandenbrouck, Marcus Geelnard, and Intel. *RcppParallel: Parallel Programming Tools for 'Rcpp'*, 2019. R package version 4.4.4.
